# Mega-Enhancers Compartmentalize Transcriptionally Active Long Genes in the Brain

**DOI:** 10.1101/2023.07.19.549737

**Authors:** Ziyu Zhao, Omar A. Payán Parra, Shin-ichiro Fujita, Francesco Musella, Nicolas Scrutton Alvarado, Frank Alber, Tomoko Yamada, Yue Yang

## Abstract

Exceptionally long genes and *cis*-regulatory enhancers are selectively activated in mammalian brain neurons, and these loci are mutation hotspots in neurological disorders. However, the organization of these large genomic elements at the level of chromosome folding, beyond local enhancer-promoter interactions, remains poorly understood. Here, we report the discovery of a nuclear subcompartment in the mouse cerebellum formed by near-megabase long enhancers and their associated long genes encoding synaptic or signaling proteins. Genomic regions within this subcompartment reside in the outer-half of the nucleus, separated from other transcriptionally active structures. Using an *in vivo* CRISPR genetic mini-screen, we uncover a specific role for the transcription factor Etv1 in coupling the compartmentalization of neuronal long genes with their expression. Together, our study defines mechanisms that organize transcriptionally active genes across chromosomes in the mammalian brain.

## INTRODUCTION

Transcriptional programs drive neuronal specification and their integration into neural circuits^1–3^. Transcription factors and chromatin enzymes control these gene expression programs by recruiting transcription machinery to gene promoters and enhancers^4–6^. Recent advances in chromatin biology have also revealed important roles for the three-dimensional (3D) organization of the genome in coordinating developmental programs of gene expression, which are specific to cell types^7–11^.

Neurons in the mammalian brain express a large group of exceptionally long genes, each spanning up to several megabases of DNA^12^. These neuronal long genes encode synaptic and transmembrane proteins, which help establish brain circuit architectures that support cognitive and sensorimotor functions in animals^12, 13^. With the evolution of long genes in vertebrates, there is a concomitant genome-wide expansion of *cis*-regulatory enhancer elements, which are selectively active in neurons and often found along the bodies of long genes^14, 15^. The dysregulation of long genes and their regulatory enhancers are commonly observed in neurological disorders of the developing and aging human brain, revealing their essential roles in cognition and behavior^13, 16–19^. These enhancers regulate transcription by forming genomic interactions with gene promoters, typically within topologically associated domains (TADs) that span up to 2-3Mb^20–24^. However, whether and how neuronal long genes and their enhancers are organized over genomic scales beyond TADs to orchestrate gene transcription in the mammalian brain is poorly understood.

At the chromosomal scale, the genome is generally partitioned into a transcriptionally active A compartment and a repressive B compartment. In mammalian cells, the A compartment partly reflects the recruitment of active genes to nuclear speckles, which are nuclear bodies enriched with RNA splicing machinery^25^. Nuclear speckles are often found in the nuclear interior, localized away from heterochromatin regions anchored to the nuclear lamina^26^. Besides nuclear speckles and lamina-associated heterochromatin, recent studies have revealed additional transcriptionally active and repressive nuclear compartments^10, 20, 26–30^. Compartments may be unique to cell types including neurons in the mammalian brain, but the molecular mechanisms that produce these structures remain a mystery.

As the most abundant neurons in the brain, cerebellar granule neurons offer a powerful system for investigating the mechanisms that orchestrate genome organization and gene transcription during development and in adulthood^22, 31, 32^. Notably, we and others have recently uncovered the presence of long-distance genomic interactions that span beyond TADs in the mouse and human cerebellum^22, 33^. In this study, we provide new insights into the microenvironments and mechanisms that generate these higher-order genomic interactions. Through deep genome architecture profiling, we identified a large group of ultra-long-distance interactions that, during development, assemble into a distinct nuclear subcompartment enriched for near-megabase-long regulatory enhancers and their associated neuronal long genes. Using split-pool recognition of interactions by tag extension (SPRITE), data-driven genome modeling, and *in situ* imaging approaches, we discovered that the enhancer-dense subcompartment occupied a microenvironment in the outer-half of the nucleus, separated from other transcriptionally active subcompartments. We next performed an *in vivo* CRISPR genetic mini-screen and found that the transcription factor Etv1 recruits its target neuronal long genes to the enhancer-dense subcompartment. Moreover, the association of Etv1 with its target enhancers promotes compartmentalization, while this process does not require gene transcription. Collectively, our findings define a novel nuclear subcompartment that drives the higher-order 3D organization of neuronal genes in the mammalian brain.

## RESULTS

### Distinct transcriptionally active subcompartments form during mouse cerebellar development

To elucidate the principles of higher-order genome organization in the mammalian brain, we employed *in situ* chromosome conformation capture with high-throughput sequencing (Hi-C) to profile the mouse cerebellum, a key brain area necessary for motor behavior and cognition^34, 35^. Our ultra-deep genome architectural profiles in this study totaled 31.7 billion reads including 12.6 billion uniquely mapped contacts (Supplementary Table 1), providing detailed maps of this brain area.

We first compared two developmental phases: at postnatal day 6 (P6), when the majority of the cerebellum comprises granule cell precursors and immature postmitotic granule neurons, and at postnatal day 22 (P22), when granule neurons have matured and become integrated into neural circuits (Fig. 1a). Hi-C analyses revealed a dramatic increase in ultra-long-distance intra-chromosomal interactions (log_2_ observed/expected>2.5; Methods), spanning 3Mb to 200Mb, during cerebellar development (Fig. 1b and Extended Data Fig. 1a). We examined the regions forming these ultra-long-distance interactions and found that most were associated with the transcriptionally active A compartment (Extended Data Fig. 1b, c). We also observed that these regions formed preferential inter-chromosomal interactions, such as along regions of chromosomes 1 and 12, which is consistent with compartment-like interactions that may represent alternative conformations across cells (Fig. 1c and Extended Data Fig. 1d). Together, these results suggest that the massive numbers of ultra-long-distance interactions formed during cerebellar development occur primarily between transcriptionally active genomic loci.

**Fig. 1.**
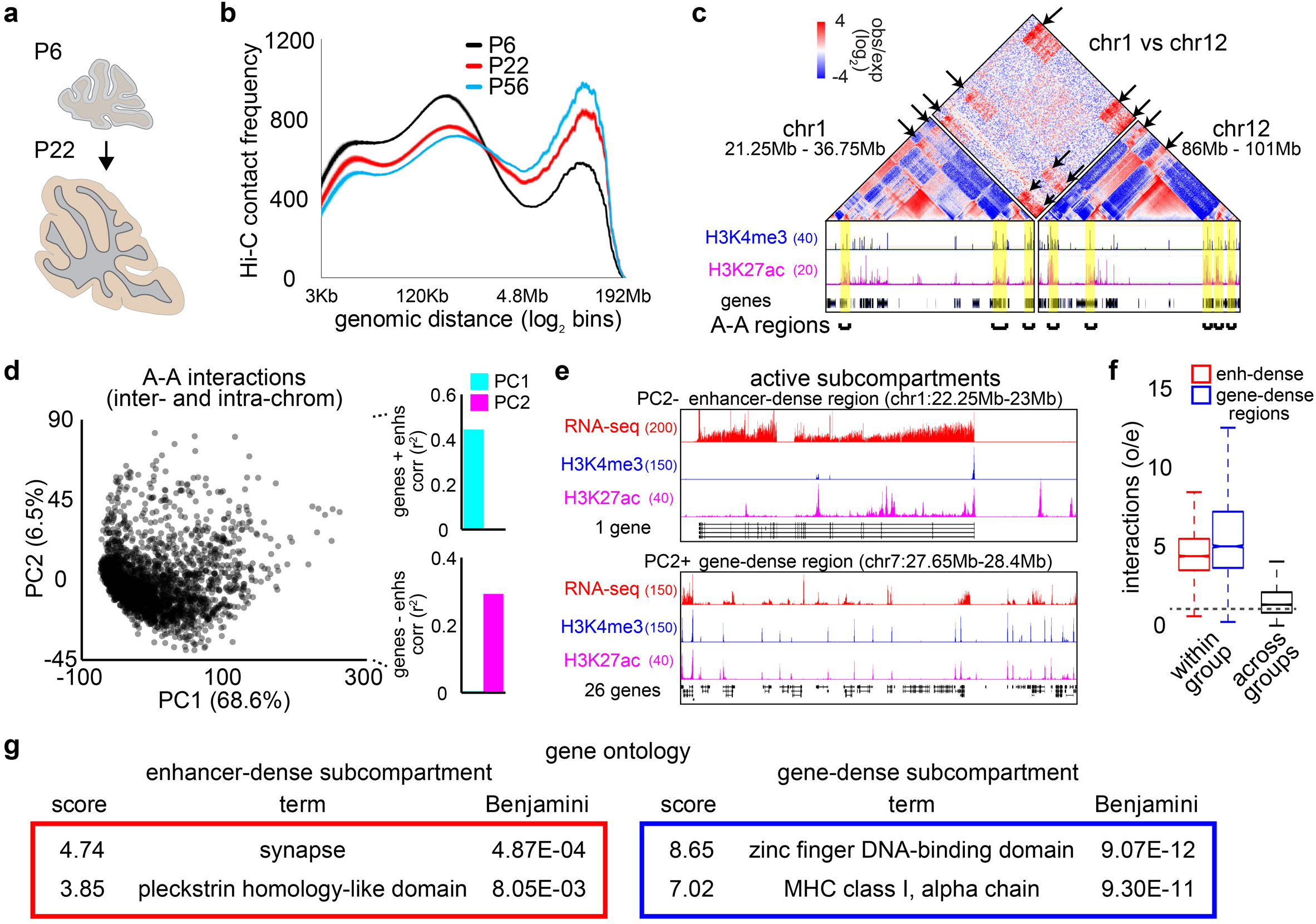
Distinct transcriptionally active subcompartments emerge in the developing cerebellum. **a**, The cerebellum from postnatal day 6 (P6) or 22 (P22) mice was dissected and subjected to Hi-C analyses. **b**, Distribution of intra-chromosomal contact frequencies at various genomic distances in the cerebellum of P6, P22, or P56^22^ mice (n = 3 biological replicates). **c**, An example of ultra-long-distance genomic interactions spanning across chromosomes 1 and 12, aligned with UCSC genome browser tracks of the active histone marks H3K4me3 and H3K27ac in P22 cerebellum. Arrows denote ultra-long-distance interactions within the A compartment and yellow highlights denote regions that form these interactions, with zoomed-in tracks in **e**, top, and Extended Data Fig. 1d. **d**, Left, principal component analysis (PCA) of regions forming ultra-long-distance interactions. Right, Pearson correlation of principal components 1 (PC1) and 2 (PC2) with various features. PC1 correlated with the density of transcriptionally active genes and enhancers. PC2 correlated with the density of enhancers minus the density of genes. **e**, UCSC genome browser tracks depicting examples of a PC2 negative (PC-) enhancer-dense region (top) and a PC2 positive (PC2+) gene-dense region (bottom), which belong to distinct active subcompartments. **f**, Interaction frequencies within and between subcompartments. Enhancer-dense regions interacted with each other but failed to interact with gene-dense regions (*P* = 0, Kruskal-Wallis test with Dunn’s post hoc test). **g**, Gene ontology analyses of active subcompartments. Enhancer-dense regions contained genes encoding for synaptic and signaling proteins. Box plots in **f** show median, quartiles (box), and range (whiskers). Data in **b** show mean and shading denote s.e.m.

We next determined additional genomic features of regions forming ultra-long-distance intra-chromosomal and inter-chromosomal interactions by performing dimensionality reduction on the interaction matrix (Extended Data Fig. 1e). Using principal component analysis (PCA), we found that the first principal component (PC1) correlated with the density of transcriptionally active genes and enhancers, which is the hallmark of the A compartment (Fig. 1d). Interestingly, the second principal component (PC2) correlated with gene density relative to enhancer density within these regions. A PC2-negative genomic region is exemplified by the *Rims1* locus, which harbors dense stretches of enhancers located within its 519Kb gene body (Fig. 1e, top). By contrast, PC2-positive regions were characterized by dense stretches of genes (Fig. 1e, bottom). Notably, this separation of regions along PC2 was also independently observed by non-negative matrix factorization (NMF; Extended Data Fig. 1e, f). These analyses reveal that a subset of regions forming extensive inter-chromosomal interactions can be separated into distinct "enhancer-dense" and "gene-dense" subcompartments, comprising 3.7% and 2.2% of the A compartment, respectively, with robust interactions occurring within each subcompartment but not between them (Fig. 1f; Methods).

The identification of distinct active subcompartments led us next to examine the genes located within them. The enhancer-dense subcompartment was enriched for genes that spanned over 100Kb and encoded synaptic and membrane-targeted signaling proteins (Fig. 1g and Extended Data Fig. 1g). On the other hand, the gene-dense subcompartment was enriched for shorter genes encoding zinc finger proteins and major histocompatibility complex proteins (Fig. 1g and Extended Data Fig. 1g). Together, these results suggest the presence of distinct active subcompartments containing different types of transcriptionally active genes in mature cerebellar granule neurons.

### Mega-enhancers emerge during development and form a nuclear subcompartment

The discovery of a distinct enhancer-dense subcompartment enriched for neuronal long genes in the cerebellum led us to further characterize the functional and structural properties of the regions it comprises. Remarkably, we found expansive H3K27ac-marked enhancer domains spanning hundreds of kilobases that emerged during cerebellar maturation, with the longest of these regions reaching over 500Kb in the adult cerebellum (Fig. 2a and Extended Data Fig. 2a). Thus, we termed these enhancer domains spanning over 100Kb as “mega-enhancers” due to their considerably greater length compared to super-enhancers, which have been reported to span ∼10Kb in proliferating cells^36^. Similar observations were found in humans, where the brain harbored some of the longest active enhancer domains compared to other tissues (Extended Data Fig. 2b)^17, 37^. We asked if mega-enhancers shared characteristics with super-enhancers, such as a close association with genes that define cell identity^36^. Genes overlapping with active mega- enhancers in granule neurons showed higher levels of expression in these neurons compared to other cerebellar cell types (Extended Data Fig. 2c), suggesting that mega-enhancers are associated with the expression of cell identity genes. In contrast, genes within the gene-dense subcompartment exhibited broad expression across various cell types in the cerebellum (Extended Data Fig. 2c). Moreover, this subcompartment was enriched for housekeeping or metabolic genes, which are known to cluster together via promoter-promoter interactions in mitotic cells (Extended Data Fig. 2d)^38^.

**Fig. 2.**
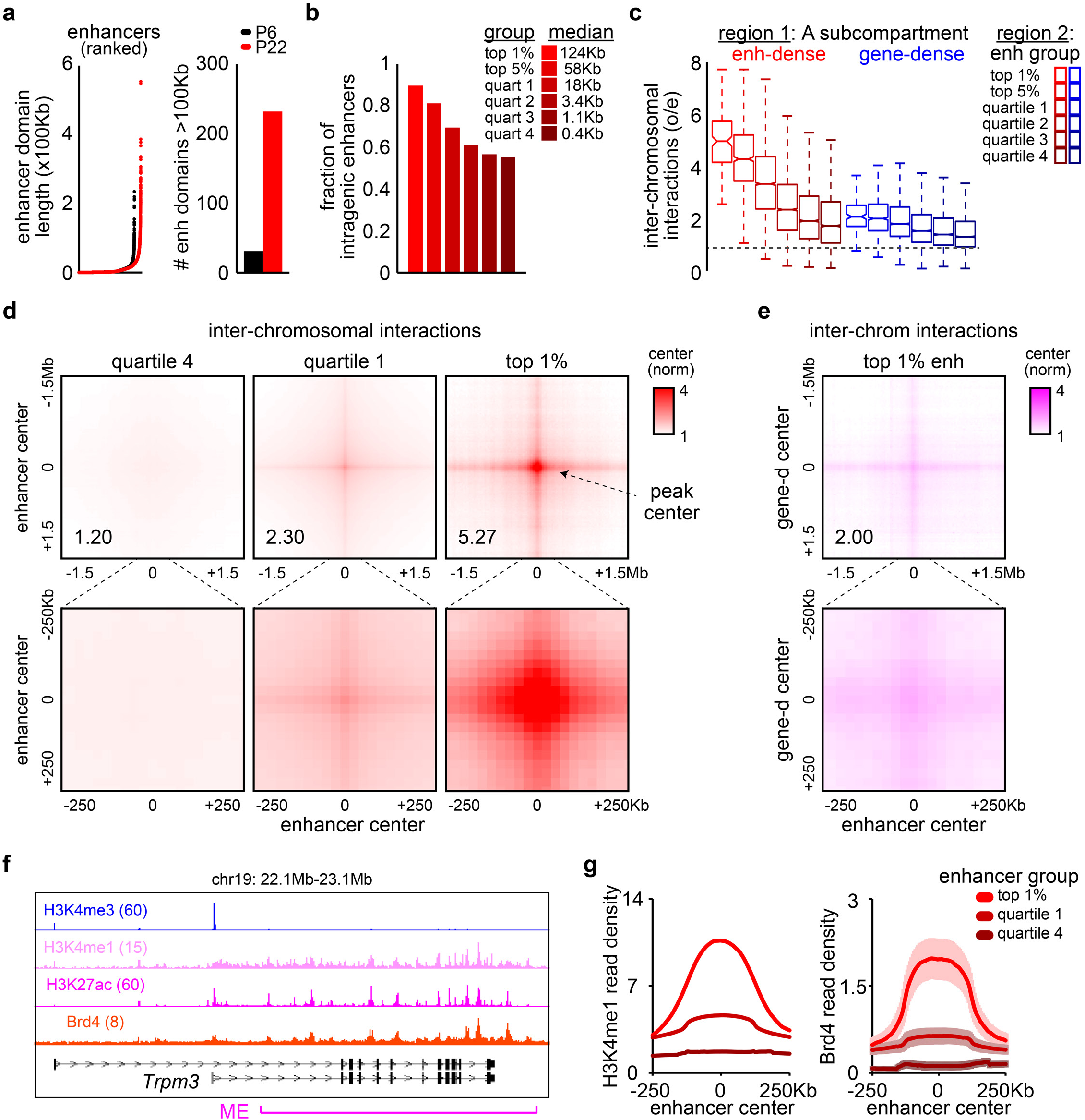
Mega-enhancers organize into a nuclear subcompartment in mature cerebellar granule neurons. **a**, Enhancer domains sorted by length (left) and the number of enhancer domains spanning over 100Kb (right) in postnatal day 6 or 22 cerebellum. A robust increase in enhancer domains spanning over 100Kb was observed during cerebellar development. **b**, Fraction of enhancer domains overlapping with gene bodies, categorized by H3K27ac levels at enhancers (*FDR* < 0.001, top 1% or top 5% compared to the quartile 4 group, pairwise Fisher’s exact test with Benjamini-Hochberg correction). The median length of enhancer domains for each group is indicated. **c**, Inter-chromosomal interaction frequency of enhancer domains, grouped as in **b**, with the enhancer-dense or gene-dense subcompartment. The top 1% enhancers, which encompass mega-enhancers, formed significantly stronger interactions with regions within the enhancer-dense subcompartment compared to those within the gene-dense subcompartment (*FDR* < 0.05, Wilcoxon signed-rank test with Benjamini-Hochberg correction, *n* = 177, 906 enhancers for top 1%, top 5%). **d**, Aggregate peak analyses^23^ of inter-chromosomal interactions among enhancer groups as in **b**. Robust and broad inter-chromosomal interactions were formed between the top 1% enhancers. **e**, Aggregate peak analyses of inter-chromosomal interactions between the top 1% enhancers and gene-dense regions, which show modest interaction levels similar to those typically observed within the A compartment. **f**, UCSC genome browser tracks showing H3K4me3, H3K4me1, H3K27ac, and Brd4 levels at a mega-enhancer (ME) region in the *Trpm3* locus. **g**, The enrichment of H3K4me1 and Brd4 at enhancers grouped by H3K27ac levels as in **b**. Box plots in **c** show median, quartiles (box), and range (whiskers). Data in **g** show mean and shading or error bars denote s.e.m.

We next examined the relationship between the activity of enhancers and the strength of their interactions with the enhancer-dense subcompartment. We categorized enhancers based on H3K27ac-enrichment, which reflects enhancer activity^36^, and analyzed their inter-chromosomal interactions with the enhancer-dense subcompartment (Fig. 2b, c and Extended Data Fig. 2e). The top 1% ranked enhancers, which include mega-enhancers spanning over 100Kb, were enriched within gene bodies and exhibited robust interactions with the enhancer-dense subcompartment, but not with the gene-dense subcompartment (Fig. 2b-2f). However, enhancers in quartile 1, with a median span of ∼18Kb and comparable in size to super-enhancers found in non-neuronal cells, only exhibited mild interactions with the enhancer-dense subcompartment (Fig. 2c, d). We also found that mega-enhancers were highly enriched for the transcriptional coactivator Brd4, a protein that regulates the expression of cell identity genes, and H3K4me1, which marks both active and primed enhancers (Fig. 2f, g)^39, 40^. These results suggest that transcriptionally active mega- enhancers form inter-chromosomal hubs in neurons.

### Transcriptionally active nuclear subcompartments occupy distinct microenvironments in neurons

We next characterized the higher-order assembly of genomic loci within the enhancer-dense subcompartment. However, a limitation of the Hi-C technique is that it captures pairwise interactions between genomic loci within close spatial proximity^41^. Therefore, we took advantage of the split-pool recognition of interactions by tag extension (SPRITE) approach, which allows for the interrogation of simultaneously interacting genomic loci within DNA-protein complexes of varying sizes (Fig. 3a)^25^. SPRITE contact maps of P22 cerebellum showed increased resolution of long-distance genomic interactions compared to Hi-C analyses (Fig. 3b), which has been also observed in other cell types^25^.

**Fig. 3.**
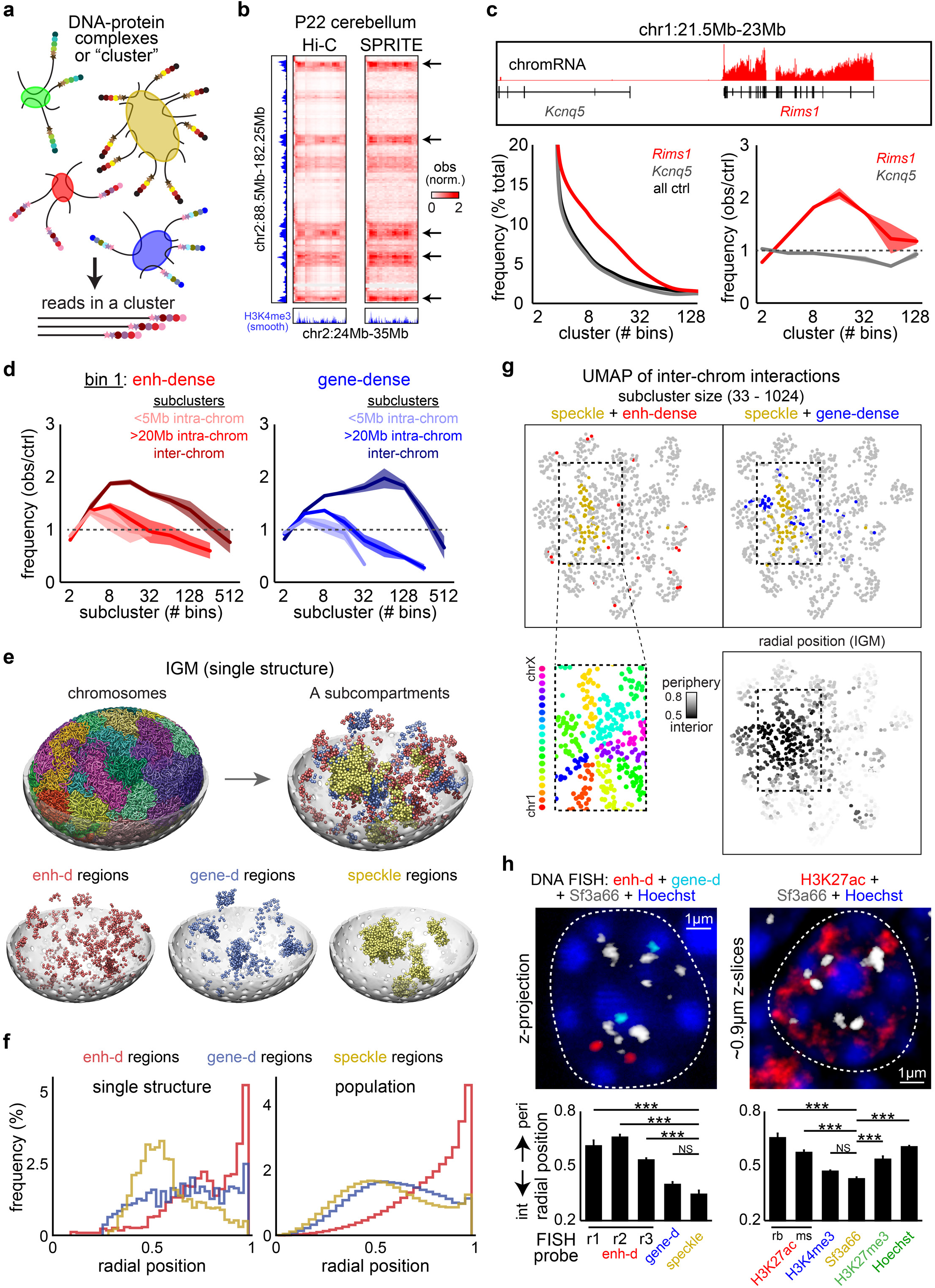
Mega-enhancers form small clusters within a distinct nuclear environment. **a**, Schematic of the SPRITE technique. Each DNA-protein complex was labeled with a unique barcode, and DNA clusters of various sizes were identified from sequencing. **b**, Long-distance interactions between regions in chromosome 2 from Hi-C (middle) or SPRITE (right) analyses together with H3K4me3 ChIP-seq tracks (left and bottom) of P22 cerebellum. Pairwise contacts from SPRITE were downweighted based on cluster size^25^. Arrows indicate ultra-long-distance genomic interactions from Hi-C. **c**, Top, UCSC genome browser track of chromatin mRNA (chromRNA) from the adult mouse cerebellum^22^. *Rims1* mRNA was highly transcribed, while little or no *Kcnq5* mRNA was detected in the adult cerebellum. Bottom left, the relative observed frequencies of the *Rims1* locus, the *Kcnq5* locus, and control (ctrl) regions within SPRITE clusters of various sizes. Bottom right, the observed frequencies of the *Rims1* or *Kcnq5* locus within SPRITE clusters normalized to that of control regions. The *Rims1* locus was observed at higher frequencies within SPRITE clusters of sizes 4 to 32 compared to control regions (*n* = 2 biological replicates). **d**, Normalized frequencies of enhancer-dense (left) or gene-dense (right) subcompartment regions appearing in SPRITE subclusters. Subclusters were selected based on the genomic distances of loci from either the same chromosome (<5MB or >20Mb, intra-chrom) or from different chromosomes (inter-chrom) in relation to enhancer-dense or gene-dense subcompartment regions. **e**, Top, the structure of an individual granule neuron nucleus generated with IGM. Bottom, IGM models highlighting the position of enhancer-dense (red), gene-dense (blue), and nuclear speckle-associated (yellow) regions, showing their center of mass. **f**, IGM predicted radial distributions of enhancer-dense (red), gene-dense (blue), and nuclear speckle- associated (yellow) regions. A single structure (left) and population average (right) are shown. **g**, UMAP plot representing the inter-chromosomal interaction preferences of regions forming ultra-long-distance Hi-C interactions within large SPRITE subclusters. Top, enhancer-dense (red), gene-dense (blue), or nuclear speckle-associated regions (gold) are highlighted. Bottom left, the chromosomes that form a distinct grouping on the UMAP plots from the top panels are enriched for genomic loci associated with nuclear speckles and the gene-dense subcompartment. Bottom right, radial positions of regions calculated from IGM and projected onto the UMAP plot of large SPRITE subclusters. **h**, Top, examples of granule neurons from adult cerebellum labeled using the Sf3a66 antibody (white) and the Hoechst DNA dye (blue), together with DNA FISH probes targeting genomic loci within the enhancer-dense (red) and gene-dense (cyan) subcompartments (left) or with the H3K27ac antibody (red, right). The dashed lines indicate the boundaries of nuclei. Bottom, radial positions of genomic loci and nuclear markers. DNA FISH probes targeting *Rims1* (r1), *Plcb4* (r2), and *Ptprt* (r3) loci were used to identify regions within the enhancer-dense subcompartment. H3K27ac antibodies were sourced from rabbit (rb) and mouse (ms). Genomic loci and nuclear markers associated with the enhancer-dense subcompartment were positioned radially away from nuclear speckles (\*\*\**P* < 0.001, one-way ANOVA with Tukey’s post hoc test, *n* = 3 biological replicates). Data in **c**, **d**, and **h** show mean and error bars or shading denote s.e.m.

To assess how genomic loci were organized within nuclear subcompartments using SPRITE, we divided the genome into 500Kb bins and calculated the number of unique bins found within each DNA cluster. We also stratified SPRITE clusters based on the number of bins to determine how genome organization relates to the size of DNA-protein complexes. Interestingly, the *Rims1* locus, which belongs to the enhancer-dense subcompartment, formed multi-way interactions within small SPRITE clusters at frequencies above those formed by control genomic regions that were not involved in long-distance interactions from Hi-C analyses (Fig. 3c). We further divided these clusters based on the distance between a genomic locus and other bins within the cluster. We found that the *Rims1* locus was enriched in subclusters where other bins were over 20Mb away or located on different chromosomes, suggesting that *Rims1* formed multi-way interactions over long genomic distances (Extended Data Fig. 3a). Extending these analyses to all regions within active subcompartments, we found that the enhancer-dense and gene-dense subcompartments each engaged in inter-chromosomal interactions above control levels (Fig. 3d). Notably, the enhancer-dense subcompartment was enriched in smaller SPRITE subclusters compared to the gene-dense subcompartment (Fig. 3d). We made similar observations with unsupervised clustering using Uniform Manifold Approximation and Projection (UMAP), where within smaller SPRITE clusters, a group of genomic loci sharing similar inter-chromosomal interaction preferences were enriched for the enhancer-dense subcompartment (Extended Data Fig. 3b, c). These results suggest that genomic regions within the enhancer-dense subcompartment engage in multi-way inter-chromosomal interactions within small genomic clusters.

Our insights into the higher-order assembly of regions within transcriptionally active subcompartments in granule neurons led us to further investigate how these subcompartments might be positioned within the nuclear topography, including their relationship with several nuclear landmarks. We used our integrative genome modeling platform (IGM)^42, 43^ to generate a population of single cell 3D genome structures of granule neurons that were statistically consistent with the Hi-C datasets obtained from P22 cerebellum (Fig. 3e). We previously demonstrated that such structures can effectively predict the folding of chromosomes relative to nuclear speckles, nucleoli, and the lamina compartments, exhibiting good agreement with experimental measurements^42, 43^. We first studied the radial positions of chromatin within the nucleus. Interestingly, we observed that enhancer-dense regions were primarily located in the outer-half of the nucleus, in contrast to the more interior locations of gene-dense regions (Fig. 3e, f). We compared these interactions with those of active chromatin predominantly associated with nuclear speckles, as previously defined in mouse cells^25^. Genomic loci associated with nuclear speckles showed the most interior nuclear locations in our models, well separated from the enhancer-dense subcompartment. Furthermore, in our single cell models, each active subcompartment was partitioned into several distinct highly-connected spatial clusters, with the enhancer-dense subcompartment segregating into the smallest spatial clusters among all active subcompartments (Fig. 3e). We next characterized each genomic locus using 14 topographic markers, which collectively describe its nuclear microenvironment by specifying its position and association frequencies to nuclear speckles, nucleoli, and the lamina as well as local chromatin compaction (Extended Data Fig. 3d)^42, 43^. Enhancer-dense regions showed distinct enrichment patterns of these topographic markers in comparison to gene-dense and nuclear speckle-associated regions (Extended Data Fig. 3d, e). As a result, all three active subcompartments occupy distinct regions in the t-SNE projections of their topographic feature vectors (Extended Data Fig. 3f), thus confirming the spatial segregation of these active subcompartments.

We next incorporated the models from IGM into our SPRITE analyses to assess the relationship between radial positioning and higher-order genome organization. We focused on larger SPRITE clusters, which better represented global genome structure and the interactions between different subcompartments. We observed with UMAP that the gene-dense subcompartment, nuclear speckles, and other genomic loci predicted to localize in the nuclear interior were partially overlapping (Fig. 3g). Conversely, the enhancer-dense subcompartment, which is predicted to be peripherally located, was found to be distributed away from the gene-dense subcompartment and nuclear speckles in the UMAP visualization of large SPRITE clusters (Fig. 3g). To test the predictions of the nuclear localization of subcompartments from IGM and SPRITE analyses, we subsequently performed DNA FISH and immunohistochemical analyses to visualize the enhancer-dense subcompartment, gene-dense subcompartment, and nuclear speckles in granule neurons in the adult cerebellum. The radial positions of genomic loci measured with imaging aligned with the predictions made by IGM modeling and confirmed that the enhancer-dense subcompartment was localized to the outer-half of the nucleus in granule neurons (Fig. 3h and Extended Data Fig. 3g, h). We also observed outward nuclear localization for the H3K27ac mark, which was associated with mega-enhancers (Fig. 3h and Extended Data Fig. 3g). In contrast, the inner half of the nucleus was concentrated for genomic loci from the gene-dense subcompartment and nuclear speckles, along with their associated protein markers (Fig. 3h and Extended Data Fig. 3g, h). Together, our findings show that transcriptionally active chromatin in granule neurons is organized into distinct subcompartments, each characterized by a distinct nuclear microenvironment.

### Dynamic changes in genome architecture during different stages of granule neuron development

The identification of a more peripheral nuclear localization for the enhancer-dense subcompartment in granule neurons led us next to determine how these genomic structures are established as granule neurons differentiate and integrate into cerebellar circuits. To study granule neurons at specific developmental stages, we used *in vivo* electroporation to label with the fluorescent protein mCherry a pool of immature granule neurons that synchronously undergo differentiation *in vivo* (Fig. 4a)^31^. At certain timepoints after electroporation, stereotypical changes in dendrite morphogenesis were observed^31^, including the formation of a leading edge process during granule neuron migration at day 2, the exuberant growth of dendrites at day 4, the pruning of excess dendrites and formation of immature postsynaptic structures at day 8, and the maturation of dendritic claw synapses at days 14 and 28 (Fig. 4a). The gene expression programs for these synchronously differentiating granule neurons were determined using translating ribosome affinity purification and sequencing (sync-TRAP-seq), and visualized on a UMAP plot of snRNA-seq data (Fig. 4b and Extended Data Fig. 4a; Supplementary Table 2)^31, 44^.

**Fig. 4.**
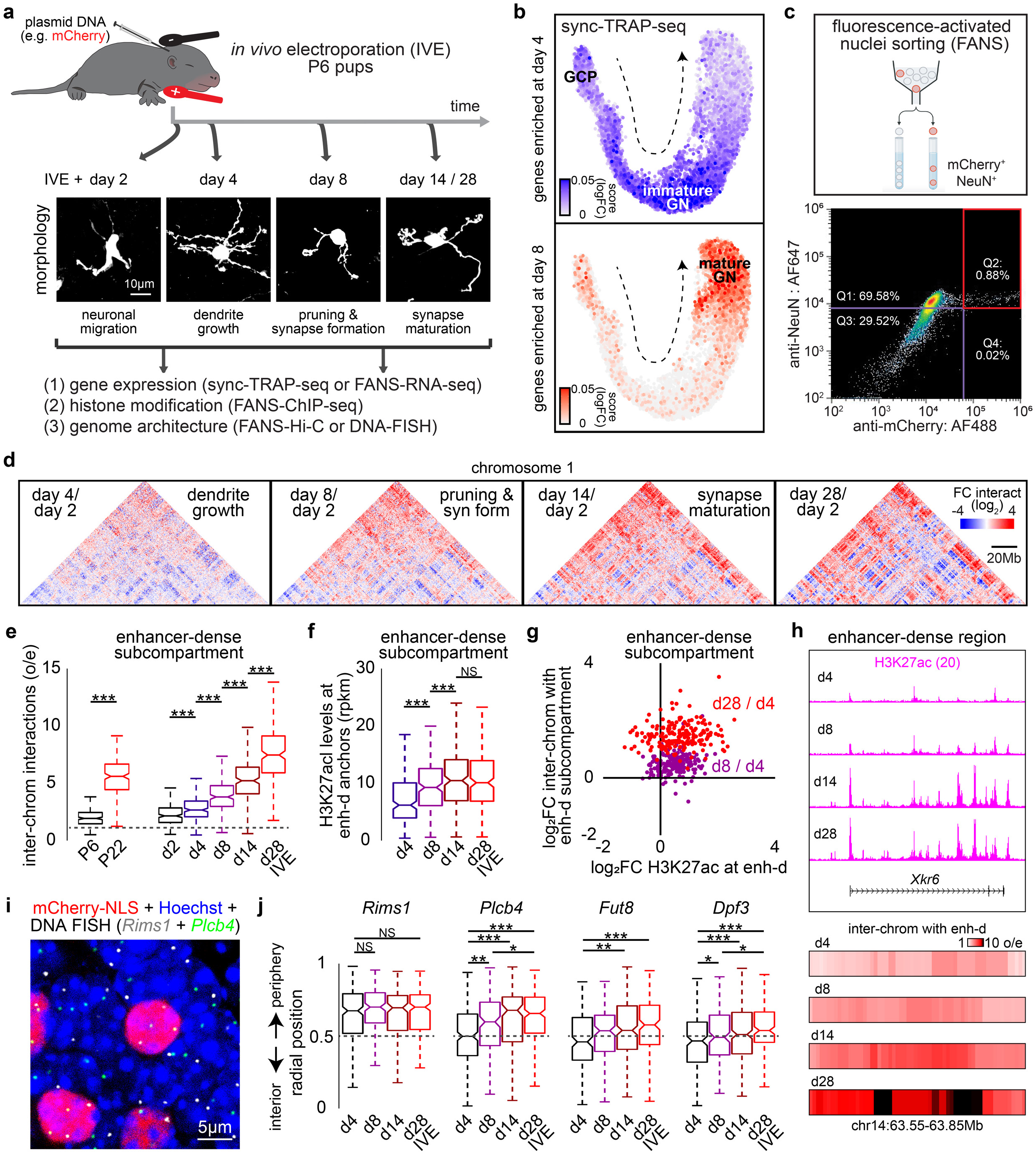
Dynamic reorganization of the enhancer-dense subcompartment across stages of granule neuron maturation. **a**, Design of the *in vivo* electroporation (IVE) approach followed by morphological analyses or genomic profiling of synchronously differentiating granule neurons at various stages of development. The dendrites of granule neurons undergo stereotyped morphological changes during the periods of granule neuron migration, dendrite growth and pruning, and dendritic claw synapse formation and maturation^31^. **b**, Gene expression profiles of synchronously differentiating granule neurons using translating ribosomal affinity purification followed by sequencing (sync-TRAP-seq). Genes enriched in granule neurons at day 4 or 8 after electroporation were visualized using a UMAP plot of snRNA-seq data from P11 cerebellum^44^. The arrow indicates the developmental trajectory of granule cell precursors (GCP) and granule neurons (GN) at various stages of differentiation. **c**, FANS procedure for isolating granule neurons electroporated with the mCherry-NLS plasmid using antibodies targeting mCherry and the granule neuron-specific marker NeuN (top), and fluorescence intensities of labeled cells and percentages of sorted positive cells with the gate Q2 (bottom, red outline). **d**, Fold changes in genomic interactions on chromosome 1 during postmitotic granule neuron maturation *in vivo*. **e**, The strength of inter-chromosomal interactions between regions within the enhancer-dense subcompartment increased during cerebellar development (left, \*\*\**P* < 0.001, Wilcoxon signed-rank test, *n* = 195 loci) or during various stages of granule neuron maturation *in vivo* (right, \*\*\**P* < 0.001, Friedman test with Nemenyi’s post hoc test, *n* = 192 loci). **f**, H3K27ac levels at enhancers within enhancer-dense regions increased from day 4 to day 14 after electroporation, then remained stable through day 28 (\*\*\**P* < 0.001, Friedman test with Nemenyi’s post hoc test, *n* = 191 loci). **g**, Changes in H3K27ac levels versus changes in inter-chromosomal interactions for enhancer-dense regions across granule neuron development. At later developmental stages, increases in compartmentalization were larger than changes in H3K27ac levels. **h**, Tracks showing H3K27ac levels at the *Xkr6* locus (top) aligned to Hi-C plots depicting the aggregate inter-chromosomal interactions between the *Xkr6* locus and the enhancer-dense subcompartment (bottom) at day 4, 8, 14, or 28 after electroporation. **i,** Images of granule neurons electroporated with the mCherry-NLS expression plasmid and labeled with the mCherry antibody (red) together with DNA FISH probes targeting the *Rims1* (grey) locus and the *Plcb4* (green) locus and with the Hoechst DNA dye (blue). **j**, Radial positioning of genomic loci within the enhancer-dense subcompartment during granule neuron maturation *in vivo*. A subset of regions within the enhancer-dense subcompartment moved toward the nuclear periphery during development (\**P* < 0.05, ** *P* < 0.01, \*\*\**P* < 0.001, Kruskal-Wallis test with Dunn’s post hoc test, *n* = 144-235 loci). Box plots in **e**, **f**, and **j** show median, quartiles (box), and range (whiskers).

We next examined the organization of the enhancer-dense subcompartment and other active genomic regions across granule neuron development by performing fluorescence-activated nuclei sorting (FANS) followed by Hi-C analyses at day 2, 4, 8, 14 or 28 after electroporation (Fig. 4c-e and Extended Data Fig. 4b-e). Remarkably, the enhancer-dense subcompartment displayed progressive increases in inter-chromosomal interaction frequency as granule neurons matured, with more pronounced developmental changes than the gene-dense subcompartment (Fig. 4e and Extended Data Fig. 4e). To determine whether changes in enhancer activity accompanied this compartmentalization, we performed FANS-ChIP-seq analyses using the H3K27ac antibody over this developmental period (Fig. 4f-h). H3K27ac levels within enhancer-dense regions increased until day 14 and were maintained thereafter, while compartmentalization continued to strengthen beyond day 14 (Fig. 4f-h). These results reveal the dynamic reorganization of inter-chromosomal interactions and enhancer activity throughout granule neuron development.

Because the enhancer-dense subcompartment maintains a unique locale in the outer-half of the nucleus in mature granule neurons, we investigated whether the positioning of these regions may be regulated during development. We analyzed granule neurons at day 4, 8, 14, or 28 after electroporation by performing mCherry immunolabeling combined with DNA FISH using probes targeting several neuronal long genes located within the enhancer-dense subcompartment (Fig. 4i, j). Interestingly, the long genes *Dpf3*, *Fut8*, and *Plcb4* moved toward the nuclear periphery as granule neurons matured *in vivo*, while the *Rims1* gene was already positioned in the outer-half of the nucleus in immature granule neurons (Fig. 4j). Besides their developmental regulation, the radial positioning of these genes was also associated with the density of CpG islands across surrounding genomic regions (Extended Data Fig. 4f-i), as found in other cell types^27, 45^. However, the developmental activation of mega-enhancers was associated with the re-localization of these neuronal long genes away from neighboring CpG-rich regions in granule neurons.

### *In vivo* CRISPR mini-screen identifies regulators of active subcompartments in granule neurons

We next applied our *in vivo* genetic approach to investigate the molecular mechanisms that regulate the enhancer-dense subcompartment in developing granule neurons. We took advantage of the CRISPR knockdown or knockout technique to target known regulators of TADs, including the cohesin subunit Rad21 and the cohesin releasing factor Wapl; structural proteins at the nuclear envelope, including Lamin A/C and the nuclear pore complex subunit Nup153; and regulators of the topological state of genomic DNA, including Top1 and Top2b. We also targeted factors recruited to enhancers or promoter-proximal regions, including the histone acetylation readers Brd4 and Brd2, the mediator complex subunit Med1, the histone acetyltransferase p300, and the transcription factor Etv1, which is associated with granule neuron maturation^46^. To induce depletion of these targets in the mouse cerebellum, we used transgenic mice expressing dCas9- KRAB or mice conditionally expressing wild type Cas9 in granule neurons, and electroporated P6 pups with the mCherry expression plasmid together with a vector encoding sgRNAs that targeted our genes of interest or control sequences (Extended Data Fig. 5a). We focused our perturbation analyses of granule neurons on day 8, 14, or 28 after electroporation and validated the efficiency of sgRNA-mediated depletion of targeted genes using RNA-seq analyses (Extended Data Fig. 5b). To assess the effects of knockdown or conditional knockout of targeted genes on genome architecture, we performed FANS-Hi-C analyses on genetically-perturbed or control granule neurons. We first conducted a joint analysis of our collection of 95 Hi-C datasets, including a small subset that we have previously generated^22, 47, 48^. We performed PCA on genome-wide A/B compartment scores across datasets and observed a trajectory of A/B compartment changes associated with the developmental maturation of granule neurons (Fig. 5a). We also observed that depletion of known structural regulators of the genome including Top2b, Wapl, or Rad21 substantially altered A/B compartments or TADs, consistent with their established functions in proliferating cells (Fig. 5b)^8, 49, 50^. Knockdown of Top2b alone, or the combined knockdown of Top1 and Top2b, weakened A compartment interactions in granule neurons, including within transcriptionally active subcompartments, and increased mixing of A/B compartments (Fig. 5b, c and Extended Data Fig. 5c-g). These effects may result from increased topological entanglements between compartments following Top2b inactivation as found in proliferating cells^50^. However, Top2b-dependent regulation of active subcompartments had little effects on neuronal long gene transcription (Extended Data Fig. 5h; Supplementary Table 3). In other experiments, we observed that conditional knockout of Rad21 selectively strengthened the enhancer-dense subcompartment, but not the gene-dense subcompartment (Fig. 5c and Extended Data Fig. 5c). Moreover, Rad21 conditional knockout reduced local intra-TAD interactions within enhancer-dense regions, suggesting that cohesin has antagonistic roles in regulating enhancer interactions within versus across TADs (Extended Data Fig. 5i-m). Together, our data indicate that known protein regulators of TADs and A/B compartments also modulate the organization of active subcompartments in granule neurons.

**Fig. 5.**
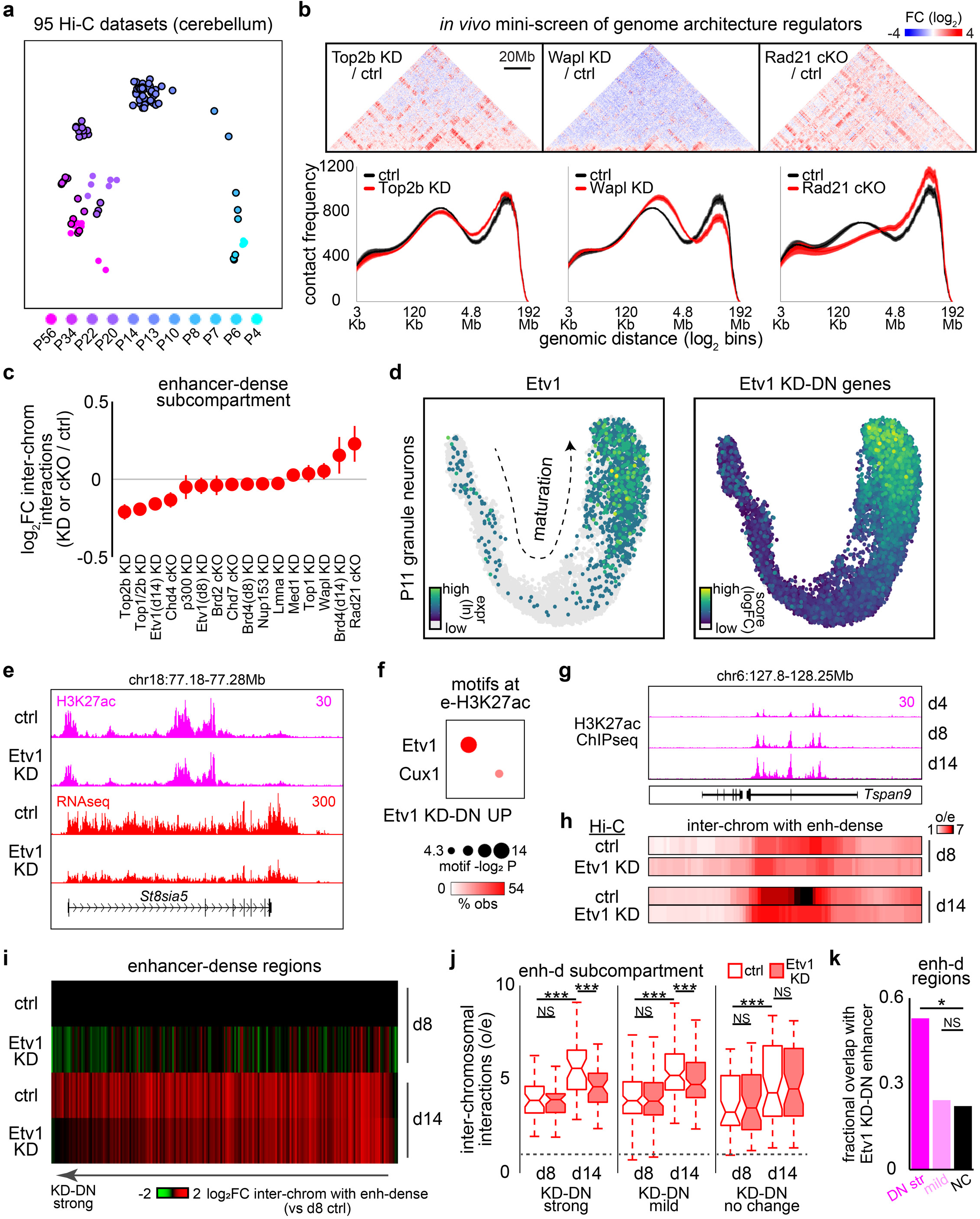
*In vivo* genetic mini-screen reveals Etv1 as a regulator of the enhancer-dense subcompartment. **a**, PCA plot of 95 Hi-C cerebellar datasets, including 73 from this study (black outline) and 22 that we have previously generated^22, 47, 48, 62^, using the A/B compartment eigenvector score derived from Juicer^63^. The first two principal components are shown. A/B compartments significantly changed throughout cerebellar development. Day 8, 14, or 28 after electroporation of P6 animals are indicated as P14, P20, or P34, respectively. **b**, Examples of targeted genes for the mini-screen for regulators of genome architecture. Granule neurons expressing the mCherry-NLS plasmid together with vectors encoding Top2b, Wapl, or control (ctrl) sgRNAs using dCas9-KRAB electroporated mice or vectors encoding Rad21 or control sgRNAs using wild type Cas9 electroporated mice were isolated with FANS and subjected to Hi-C analyses. Top, Hi-C contact maps of chromosome 1 showing the fold changes in interactions between the knockdown (KD) or conditional knockout (cKO) conditions and their respective control conditions. Bottom, histograms of intra-chromosomal contact frequencies as a function of the genomic distance between regions (*n* = 3-4 biological replicates). **c**, Fold changes in inter-chromosomal interactions between regions within the enhancer-dense subcompartment upon knockdown or conditional knockout of 13 nuclear proteins (*n* = 2-6 biological replicates). **d**, Expression of Etv1 (left) or Etv1 knockdown-downregulated genes (right) in developing granule neurons visualized with UMAP as in Fig. 4b. **e**, UCSC browser tracks of H3K27ac levels or gene expression at the *St8sia5* locus upon Etv1 knockdown in granule neurons. **f**, Transcription factor binding motifs enriched at enhancers with downregulated or upregulated H3K27ac levels upon Etv1 knockdown in granule neurons. **g**,**h**, Tracks showing H3K27ac levels at the *Tspan9* locus in granule neurons at day 4, 8, or 14 after electroporation (**g**), aligned to Hi-C plots depicting the aggregate inter-chromosomal interactions between the *Tspan9* locus and the enhancer-dense subcompartment in control or Etv1 knockdown granule neurons at day 8 or 14 after electroporation (**h**). **i**, Changes in inter-chromosomal interactions between regions within the enhancer-dense subcompartment in granule neurons at day 8 or 14 after electroporation for a control condition or after Etv1 knockdown. Interactions are normalized to those in day 8 control neurons and sorted by the fold reduction upon Etv1 knockdown compared to its control at day 14. **j**, Inter-chromosomal interactions between regions with the enhancer-dense subcompartment as in **i**, categorized based on the fold change in interactions upon Etv1 knockdown (\*\*\**P* < 0.001, Friedman test with Nemenyi’s post hoc test, *n* = 41-104 loci). **k**, Fractional overlap between the groups of genomic regions shown in **j** and enhancers with reduced H3K27ac levels upon Etv1 knockdown as in **f**. Regions exhibiting a strong dependence on Etv1 for compartmentalization (log_2_ fold change in interactions < -0.2) were significantly enriched for Etv1-regulated enhancers containing the Etv1 consensus motif (\**FDR* < 0.05, pairwise Fisher’s exact test with Benjamini-Hochberg correction, compared to NC regions). Box plots in **j** show median, quartiles (box), and range (whiskers). Data in **b** and **c** show mean and shading or error bars denote s.e.m.

### Etv1 recruits its target long genes to the enhancer-dense subcompartment

Interestingly, in our mini-screen, we discovered that the ETS variant transcription factor 1 (Etv1) promoted inter-chromosomal interactions between regions within the enhancer-dense subcompartment in granule neurons (Fig. 5c). We first characterized Etv1 expression in the cerebellum and observed that Etv1 mRNA and protein were selectively enriched in granule neurons in the internal granule layer (IGL) and upregulated during their period of synaptic development (Fig. 5d and Extended Data Fig. 6a)^46^. Consistent with these findings, RNA-seq analyses of Etv1 knockdown granule neurons at day 8 after electroporation, during their period of synaptogenesis, revealed that Etv1 was required for the expression of genes specific to mature granule neurons (Fig. 5d and Extended Data Fig. 6b; Supplementary Table 3). Changes in the expression of Etv1-regulated genes were associated with changes in H3K27ac levels at their enhancers (Fig. 5e and Extended Data Fig. 6c, d). Interrogation of DNA sequences within Etv1- regulated enhancers revealed a robust enrichment of the Etv1 consensus motif specifically at Etv1 knockdown-downregulated enhancers, but not at upregulated enhancers (Fig. 5f). These findings suggest that Etv1 activates a group of enhancers harboring its DNA-binding sequence to drive gene expression in mature granule neurons.

We next determined the relationship between the gene expression programs regulated by Etv1 and the enhancer-dense subcompartment. Interestingly, genes transcriptionally downregulated upon Etv1 knockdown were enriched within the enhancer-dense subcompartment, but not within the gene-dense subcompartment or nuclear speckle-associated regions (Extended Data Fig. 6e). An example is the Etv1 target gene *Tspan9*, which harbors a dense stretch of enhancers that became activated as granule neurons matured (Fig. 5g; Supplementary Table 3). Strikingly, the *Tspan9* locus formed increased interactions with the enhancer-dense subcompartment during granule neuron development and these genomic interactions were inhibited upon knockdown of Etv1 (Fig. 5h). These findings extended genome-wide, where loci containing Etv1 knockdown-downregulated long genes strengthened their inter-chromosomal interactions with the enhancer-dense subcompartment over development, and these changes were blocked by Etv1 knockdown (Extended Data Fig. 6f). Moreover, the effects of Etv1 knockdown on the enhancer-dense subcompartment were specific to genomic regions enriched for Etv1 knockdown-downregulated enhancers, which contain the Etv1 binding sequence (Fig. 5f, 5i-k and Extended Data Fig. 6g). These results suggest that Etv1 recruits a specific set of target long genes to the enhancer-dense subcompartment and activates their expression during granule neuron maturation.

### Compartmentalization does not require transcription but is mediated by the association of Etv1 with enhancers

To further investigate the mechanisms underlying the compartmentalization of neuronal long genes, we characterized the roles of the transcriptional coactivator Brd4, which was enriched at mega-enhancers (Fig. 2f, g). Similar to Etv1, Brd4 was required for the expression of a group of developmentally upregulated genes (Fig. 6a, b and Extended Data Fig. 7a; Supplementary Table 3). In addition, Brd4 knockdown-downregulated genes were enriched in the enhancer-dense subcompartment and strengthened their interactions with this subcompartment over development (Fig. 6c and Extended Data Fig. 7b), phenocopying our findings with Etv1-regulated genes. Surprisingly, in contrast to Etv1, Brd4 knockdown failed to reduce inter-chromosomal interactions between Brd4 knockdown-downregulated long genes and the enhancer-dense subcompartment (Fig. 6c). These findings suggest that gene transcription, which is impaired after Brd4 knockdown, is not required for establishing genomic interactions with the enhancer-dense subcompartment. Rather, these results suggest other functions of Etv1 in recruiting its target long genes to the enhancer-dense subcompartment.

**Fig. 6.**
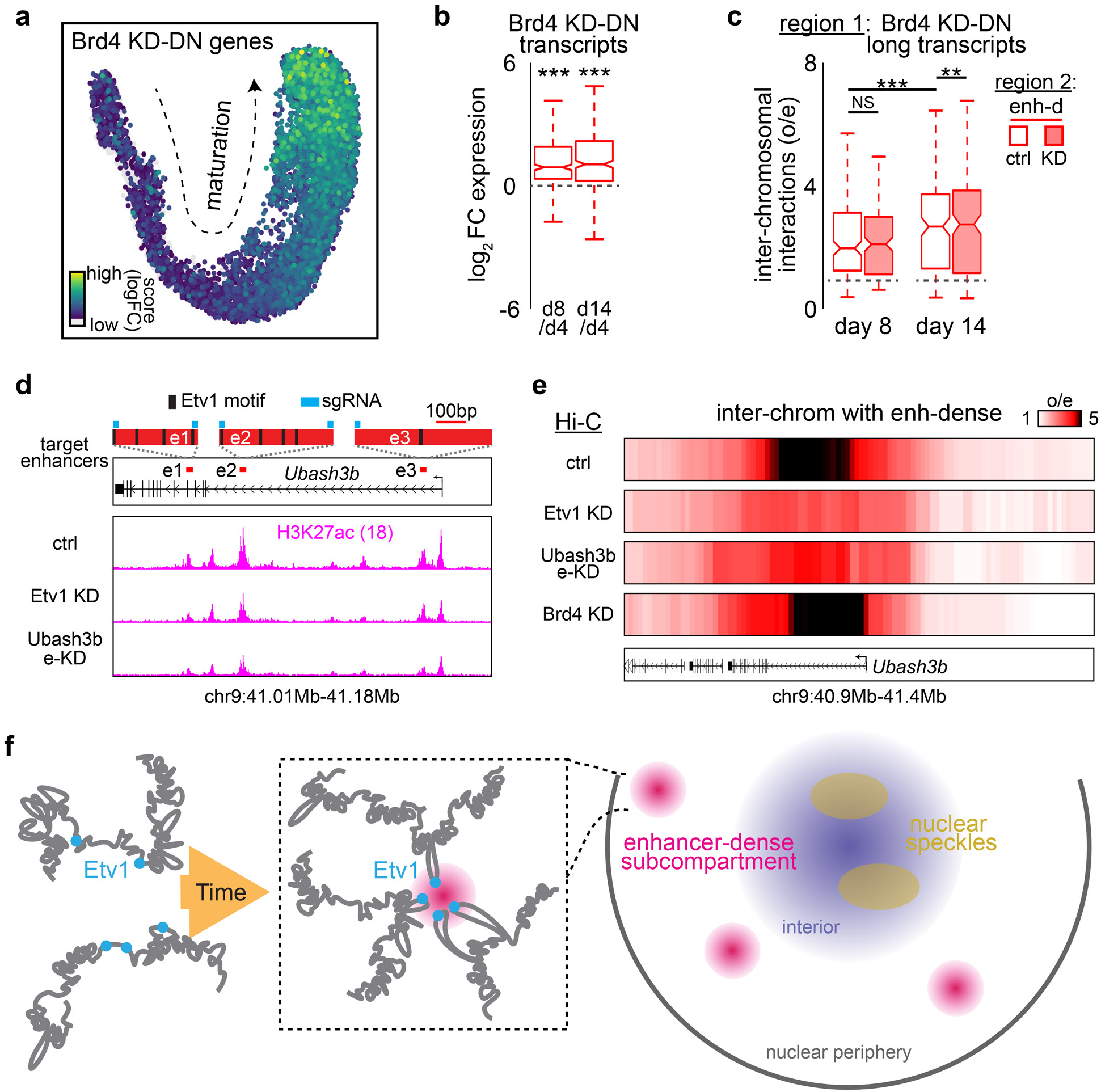
Compartmentalization occurs independently of transcription but is mediated by the association of Etv1 with enhancers. **a**, Expression of Brd4 knockdown-downregulated genes in developing granule neurons visualized with UMAP as in Fig. 4b. **b**, Changes in Brd4 knockdown-downregulated transcript expression in developing granule neurons from day 4 to day 8 or 14 after electroporation (\*\*\**P* < 0.001, Wilcoxon signed-rank test with Benjamini-Hochberg correction, *n* = 3-6 biological replicates). **c**, Hi-C analyses of inter-chromosomal interactions between Brd4 knockdown-downregulated long genes (log_2_ fold change < -0.585) and the enhancer-dense subcompartment in control or Brd4 knockdown granule neurons at day 8 or 14 after electroporation. Long genes expressed in an Brd4-dependent manner strengthened their interactions with the enhancer-dense subcompartment during granule neuron maturation *in vivo*, but Brd4 knockdown failed to block these developmental changes (\*\**P* < 0.01, \*\*\**P* < 0.001, Friedman test with Nemenyi’s post hoc test, *n* = 165 transcripts). **d**, Top, Etv1 motifs identified with JASPAR^64^ within *Ubash3b* gene enhancers. Bottom, tracks showing H3K27ac levels at the *Ubash3b* locus following Etv1 knockdown or *Ubash3b* enhancer inactivation using the dCas9-KRAB system. **e**, Hi-C contact maps depicting the aggregate inter-chromosomal interactions between the *Ubash3b* locus and the enhancer-dense subcompartment in granule neurons at day 14 after electroporation, shown for a control condition or after Etv1 knockdown, *Ubash3b* enhancer inactivation, or Brd4 knockdown. **f**, Our working model. Left, the transcription factor Etv1 (cyan) activates its target neuronal long genes and recruits them to the enhancer-dense subcompartment progressively over time as granule neurons mature. Right, the enhancer-dense subcompartment (magenta) occupies a distinct microenvironment in the outer-half of the nucleus in neurons. Nuclear speckles are depicted in gold; nuclear interior in dark blue. Box plots in **b** and **c** show median, quartiles (box), and range (whiskers).

We finally tested whether interactions between Etv1 and its target gene enhancers were required for their nuclear compartmentalization. The *Ubash3b* gene harbors multiple Etv1 consensus motifs and was sensitive to Etv1 knockdown (Fig. 6d, top). To block Etv1 interactions with *Ubash3b* gene enhancers, we targeted these enhancers with sgRNAs in dCas9-KRAB mice. In ChIP-seq analyses of electroporated granule neurons, we observed a robust reduction in H3K27ac levels at *Ubash3b* enhancers following their inactivation (Fig. 6d, bottom). Importantly, inactivating these enhancers specifically reduced their association with the enhancer-dense subcompartment, phenocopying the effects of Etv1 knockdown at the *Ubash3b* locus (Fig. 6e and Extended Data Fig. 7c). In contrast, although the transcriptional activator Brd4 also promotes Ubash3b expression, Brd4 knockdown had little or no effect on the compartmentalization of the *Ubash3b* gene (Fig. 6e; Supplementary Table 3). Together, these findings highlight the distinct functions of Etv1 in compartmentalizing its target long genes during granule neuron development (Fig. 6f).

## DISCUSSION

Our study unveils new principles underlying the higher-order genome organization of mega-enhancers and their associated neuronal long genes in the brain. We find that the prevalence of hundreds of mega-enhancers activated in mammalian brain neurons is pivotal for establishing a novel enhancer-dense subcompartment. Mega-enhancers are characterized by exceptionally long stretches of H3K27ac-marked enhancers spanning hundreds of kilobases, which are over an order of magnitude larger than super-enhancers^36^. In non-neuronal cells, super-enhancers form local interactions with neighboring gene promoters or enhancers primarily within TADs^51, 52^ and occasionally across neighboring TADs^20, 28, 51^. Certain genetic perturbations, such as the depletion of the cohesin subunit Rad21 or generation of the Brd4-NUT fusion oncogene, have been shown to induce inter-chromosomal interactions between super-enhancers or hyperacetylated TADs in proliferating cells^8, 53, 54^. Remarkably, even under physiological conditions, mega-enhancers from different chromosomes coalesce into multiple clusters within a distinct subcompartment in granule neurons.

Distinct patterns of genome compartmentalization have been observed in neuronal cell types. For example, in olfactory sensory neuron (OSN) progenitors, olfactory receptor genes are compartmentalized within heterochromatin^55^. Later, in mature OSNs, transcriptional hubs comprising a chosen olfactory receptor gene and active enhancers are positioned adjacent to the heterochromatin compartment^28, 56^. Consistent with this theme, we and others find that the gene-dense subcompartment is spatially organized alongside heterochromatin in mature neurons across the brain^20, 26, 33, 57^. However, unlike these heterochromatin-organized genome architectures, the enhancer-dense subcompartment is not primed by heterochromatin but instead emerges through the activation of hundreds of mega-enhancers across the genome. Furthermore, the extensive inter-chromosomal interactions among mega-enhancers in mammals are distinct from enhancer-promoter loops or even longer-range meta-loops, in which a gene interacts specifically with a distal regulatory element located up to megabases away^58^. Rather, the numerous interactions among mega-enhancers more closely resemble the higher-order genomic interactions organized by nuclear speckles in mammalian cells^25^.

Among the nuclear proteins tested in our mini-genetic screen, the sequence-specific transcription factor Etv1 uniquely couples the spatial compartmentalization of its target enhancer regions with gene transcription during granule neuron development *in vivo*. However, our results reveal that gene transcription is not required for forming the enhancer-dense subcompartment. Furthermore, because Etv1 selectively recruits only a subset of genes to this subcompartment, other transcription factors may similarly compartmentalize their target genes and the combined activities of these factors may collectively establish the enhancer-dense subcompartment.

The spatial distribution of nuclear bodies such as nuclear speckles or the nucleolus have well established roles in higher-order 3D genome organization and gene regulation in cells^59, 60^. By integrating experimental and computational approaches, we found that the novel enhancer-dense subcompartment is composed of smaller genomic DNA clusters. These enhancer clusters altogether form an independent microenvironment in the outer part of the nucleus. In contrast, the gene-dense subcompartment is organized by larger genomic DNA clusters concentrated in the nuclear interior alongside nuclear speckles^25^. These findings raise important questions on why mega-enhancers are positioned in a distinct locale in neurons. One possibility is that the more peripheral localization of mega-enhancers improves sensitivity to signals triggered by transmembrane receptor activation^1^. Once intracellular signals translocate through the nuclear pore, they may more readily activate mega-enhancers and their associated neuronal long genes located closer to the nuclear envelope. Alternatively, because long genes have been reported to physically expand during transcription^59^, the outer-half of the nucleus may provide additional space to accommodate the transcription of the numerous long genes found in mammalian brain neurons.

Together, our study reveals that the novel enhancer-dense subcompartment serves as the architectural backbone for organizing neuronal long genes at the chromosomal level. Because the dysregulation of neuronal long genes encoding synaptic proteins is likely causative of neurological diseases including autism spectrum disorders^13, 17, 61^, elucidating how these synaptic genes are developmentally regulated at the level of chromosomal folding will advance our understanding of how neural circuits are properly assembled in humans.

## Data availability

Raw and processed sequencing data generated for this study are available from 4DN data portal (https://data.4dnucleome.org/) deposited under Tomoko Yamada, NW.

## Acknowledgments

We thank members of the Yang Lab for helpful discussions, the Biological Imaging Facility (RRID:SCR_017767) for imaging, and Ming Hu for discussions about FIRE analyses. This project was supported by NIH 4DN grants U01DA053691 (T.Y.). and UM1HG011593 (F.A.), and R01NS123285 (Y.Y.).

## Author contributions

Z.Z., O.A.P.P, S.F., Y.Y., and T.Y. designed the study and wrote the manuscript. Z.Z., O.A.P.P, S.F., N.S.A., Y.Y. and T.Y. performed RNA-seq, ChIP-seq, Hi-C, SPRITE, DNA FISH, immunohistochemistry, *in vivo* electroporation, FANS, TRAP, and bioinformatics analyses. F.M. and F.A. generated and analyzed the IGM models and described the results.

## Author Information

The authors declare no conflicts of interest.

## Methods

### Animals

C57BL/6 or CD1 mice were purchased from Charles River Laboratories or maintained under pathogen-free conditions. Homozygous dCas9-KRAB mice (#030000, Jackson Laboratory) were bred with B6D2F1/J mice (#100006, Jackson Laboratory), or LSL-Cas9-EGFP (#026175, Jackson Laboratory) mice were bred with GABA(A)Rα6-Cre driver mice^65^ for *in vivo* electroporation. Both male and female mice were used. Mice were allocated into sex-matched, littermate-matched experimental groups. All animal experiments were done according to protocols approved by the Animal Studies Committee of Northwestern University in accordance with the National Institutes of Health guidelines. Details of the animals used for experiments are listed in Supplementary Table 4.

### Antibodies

Antibodies to H3K4me3 (Abcam, ab8580), H3K27ac (Active Motif, 39085, lot#17019020; 39134, lot#06921014; Abcam ab4729), H3K27me3 (Millipore, 07-449, lot#2919706), Sf3a66 (Abcam, ab77800), NeuN (Abcam, ab177487), Brd4 (Bethyl Laboratories, A301-985A, lot#7), H3K9me3 (Abcam, ab8898), and mCherry (Abcam, ab205402), Etv1 (Invitrogen, PA5-77975), Calbindin (Sigma, C9848) for immunohistochemistry, ChIP-seq, or FANS experiments were purchased.

### Plasmid DNA

The MLM3636 vector (addgene #43860) was modified with A-U flip and stem extension to increase the stability of sgRNA as described^66^. sgRNA was cloned into the modified MLM3636a vector using primers containing the target sequence listed in Supplementary Table 5^22^. For genes targeted with two sgRNAs, the second U6-sgRNA sequence was subcloned downstream of the first sgRNA sequence. The pCAG-Rpl10a-GFP and pCAG-mCherry vectors were previously described^31^. mCherry-NLS (addgene #58476) was subcloned into the pCAG vector to generate pCAG-mCherry-NLS.

### Statistics

Statistical analyses were performed using Matlab, Microsoft Excel, or R. Normality of data was assessed using the Shapiro–Wilk test. For experiments in which only two groups were analyzed, the t-test was used for normal distributions. For two groups with non-normal distributions, the Mann-Whitney-Wilcoxon test (unpaired data) or Wilcoxon signed-rank test (paired data) was used. When comparing specific pairs of samples among multiple groups, p-values were adjusted using the Benjamini-Hochberg procedure. Comparisons of all samples across multiple groups were performed using analysis of variance (ANOVA) followed by Tukey’s post hoc testing for normal distributions, the Kruskal-Wallis test followed by Dunn’s post hoc testing with p-values adjusted using the Benjamini-Hochberg procedure for unpaired non-normal distributions, and the Friedman test followed by Nemenyi’s post hoc testing for paired non-normal distributions. For contingency tables, the pairwise Fisher’s exact test was used to compare each condition against a control, and p-values were adjusted for multiple comparisons using the Benjamini–Hochberg procedure.

### *In vivo* electroporation

*In vivo* electroporation of CD1, dCas9-KRAB, or Gabra6-Cre; LSL-Cas9-EGFP mice conditionally expressing wild type Cas9 in granule neurons was performed as described^31, 67^. The indicated plasmids were injected into the cerebellum of P6 mouse pups and then subjected to five 50 ms electrical pulses of 135 mV with 950 ms intervals. Electroporated pups were returned to moms and subjected to biochemical or immunohistochemical analyses at the indicated days after electroporation. For morphological analyses, the cerebellum was fixed with 4% PFA in PBS and subjected to immunohistochemical analyses as described below.

### sync-TRAP

TRAP from synchronously developing granule neurons (sync-TRAP) was performed as described with modifications^31, 68, 69^. The cerebellum of P6 mouse pups was electroporated with the pCAG- Rpl10a-GFP and pCAG-mCherry plasmids together with a vector encoding U6-sgRNA targeting the indicated gene or a control sequence at day 4, 8 or 28 after electroporation. The cerebellum from mCherry-labeled mice was dissected in HHGN (1X HBSS, 2.5 mM HEPES (pH 7.4), 35 mM glucose, 4 mM sodium bicarbonate) with 100 μg/ml cycloheximide and 0.5 mM DTT under a fluorescence stereomicroscope (Olympus SZX16). Cerebellar tissue was homogenized by 10 strokes with dounce (A) followed by 10 strokes with dounce (B) in lysis buffer (50 mM Tris-HCl pH 7.4, 100 mM KCl, 12 mM MgCl_2_, 1% NP-40, 0.4 unit/ml RNase inhibitor (Promega, N2511), 1 mM DTT, 100 μg/ml cycloheximide). Lysates were incubated with 1 mg/µl heparin (Sigma, H3393) for 2 minutes on ice prior to centrifugation. Samples were spun down at 10,000*g* for 10 minutes at 4C and the supernatant was used for affinity purification of the Rpl10a-GFP and mRNA complex. Cerebellar lysates were added to Dynabeads protein G (Invitrogen) coupled with 15 µg of each monoclonal GFP antibody (HtzGFP-19F7 and HtzGFP-19C8 from Memorial Sloan Kettering Cancer Center) for a total of 30 mg. After extensive washes with washing buffer (50 mM Tris-HCl pH 7.4, 300 mM KCl, 12 mM MgCl_2_, 1% NP-40, 0.5 mM DTT, 100 μg/ml cycloheximide), immunoprecipitated RNA was purified with a RNeasy Micro Kit (Qiagen).

### FANS

The cerebellum of P6 mouse pups was electroporated with the pCAG-mCherry-NLS plasmid together with a vector encoding U6-sgRNA targeting the indicated genes or control sequences and dissected at day 1, 2, 4, 7, 8, 14, or 28 after electroporation under a stereomicroscope (Olympus SZX16) to assess electroporation efficiency. The cerebellar vermis was fixed with crosslink solution (1% PFA in PBS) by homogenizing with a glass dounce homogenizer and incubated for 10 min at room temperature. Crosslinking reaction was quenched by 125 mM glycine solution and incubated for 5 min. The cell pellet was washed twice with BSA/PBS solution (0.3 % BSA, 0.1 % Triton-X in PBS) and stored at -80C.

To isolate nuclei, the frozen pellet was thawed on ice, homogenized with lysis buffer (10 mM Tris-HCl pH 8, 10 mM NaCl, 0.2% Triton-X) and incubated on ice for 10 min. The lysate was pelleted by centrifugation at 700 *g* for 5 min at 4C and washed with lysis buffer once. The pellet was resuspended in BSA/PBS solution and filtered using a 50 µm filter (CellTrics). Filtered nuclei were incubated with antibodies against mCherry (1:1000) and NeuN (1:500) for 1 hr at 4C with rotation. Nuclei were washed with BSA/PBS solution twice and stained with secondary antibodies including anti-chicken Alexa 488 (1:250, Invitrogen, A11039) and anti-rabbit Alexa 647 (1:250, Abcam, ab150075) for 30 min at 4C with rotation, followed by washing with BSA/PBS solution twice. Stained nuclei were resuspended in BSA/PBS solution and filtered using a 50 µm filter prior to FANS. mCherry^+^/NeuN^+^ nuclei were enriched with a SH800S cell sorter (Sony) using a 100 µm nozzle sorting chip and 488/638nm lasers. Nuclei were first sorted using the ultra-yield mode and then re-sorted using the normal mode.

For RNA-seq following FANS, sorted cells were frozen and stored at -80C. ∼2,000 sorted nuclei from single or multiple animals were thawed on ice and reverse-crosslinked with 100 µL of RIP buffer (100 mM Tris-HCl pH 8.0, 10 mM EDTA, 1% SDS, 1 µl of RNAse inhibitor (Promega)) containing 1 µl of Proteinase K (New England Biolab) for 1 hr at 65C. Then, nuclear RNA was purified using a RNeasy micro kit (Qiagen) according to the manufacturer’s instructions. Half of the purified RNA was subjected to RNA-seq library preparation described below.

For Hi-C following FANS, ∼1,000-12,000 cells (Supplementary Table 1) were sorted into PBS and lysis buffer at four times the volume was added immediately. Nuclei were pelleted and resuspended with 0.5% SDS for permeabilization as described with modifications^22, 23^.

For FANS-ChIP-seq, the cerebellum of electroporated mouse pups was fixed with homogenization in 4% PFA/PBS solution for 10 min, followed by quenching with glycine solution for 5 min. After the dissociation and labeling of nuclei with mCherry and NeuN antibodies, mCherry^+^/NeuN^+^ electroporated granule neurons were enriched with FANS and subjected to sonication for DNA fragmentation. ∼40,000-45,000 sorted nuclei were used for chromatin immunoprecipitation.

### RNA-seq

Total RNA was extracted from the cerebellum of postnatal day 6 or 22 mice using Trizol (Thermo Fisher Scientific) according to the manufacturer’s instructions. ∼100 ng of RNA from the cerebellum or half the amount of purified RNA from FANS-isolated granule neurons was treated with a NEBNext rRNA Depletion Kit (New England Biolabs). RNA-seq was performed using libraries prepared with a NEBNext Ultra™ Directional RNA Library Prep Kit for Illumina (New England Biolabs).

For sync-TRAP-seq to assess knockdown or conditional knockout of target genes, libraries were prepared using the Smart-seq3 protocol^70^ with modifications. Briefly, half the amount of immunopurified RNA was subjected to reverse transcription followed by PCR amplification. 25 ng of cDNA was used for the Tn5 reaction to generate libraries. For sync-TRAP-seq to perform differential gene expression analyses, libraries were prepared using a Trio RNA-seq kit with ribo- depletion (TECAN) according to the manufacturer’s instructions.

All libraries were sequenced on the Illumina NextSeq 500 or 550 platform to obtain 36-37 bp paired-end reads. Two to eleven biological RNA-seq replicates were performed.

### ChIP-seq

ChIP-seq assays were performed as described with modifications^31, 32^. The cerebellum from postnatal day 6 or 22 mice was fixed with 1.1% formaldehyde solution. Immunoprecipitation was performed in RIPA buffer (10 mM Tris-HCl pH 8.0, 140 mM NaCl, 0.1% SDS, 1% Triton-X, 0.1% DOC, 1 mM EDTA, 0.5 mM EGTA) using the indicated antibodies with Dynabeads protein G (Thermo Fisher Scientific).

For the Brd4 antibody, Dynabeads protein G were preincubated with the antibody for several hours and washed with RIPA buffer before immunoprecipitation. Lysates were added to the antibody-coupled beads and incubated at 4C overnight. After extensive washes of beads with RIPA buffer three times, high salt RIPA buffer (10 mM Tris-HCl pH 8.0, 500 mM NaCl, 0.1% SDS, 1% Triton-X, 0.1% DOC, 1 mM EDTA, 0.5 mM EGTA) three times, LiCl buffer (10 mM Tris-HCl pH 8.0, 150 mM LiCl, 1% Triton-X, 1% DOC, 1 mM EDTA, 1 mM EGTA) and TE (10 mM Tris-HCl, 1mM EDTA) twice, tagmentation reactions were performed as described^71^. Briefly, immunoprecipitated DNA fragments on the beads were digested with Tn5 (Illumina), and free DNA fragments were washed away with RIPA buffer and TE. The DNA fragments on the beads were treated with proteinase K (Thermo Fisher Scientific) for 1 hr at 37C and de-crosslinked at 65C overnight. DNA fragments were purified with a PCR purification kit (Qiagen) and amplified with primers containing indexes for sequencing using Q5 High-Fidelity 2xMaster mix (New England Biolabs).

For the H3K27ac, H3K4me3, H3K27me3, or H3K9me3 antibody, the lysate was mixed with each antibody overnight at 4C. BSA-coated Dynabeads protein G was added and incubated at 4C for 1 hr. After extensive washes of beads with RIPA buffer three times, high salt RIPA buffer three times, and TE three times, DNA fragments were eluted in elution buffer (10 mM Tris-HCl pH 8.0, 350mM NaCl, 1% SDS, 0.1 mM EDTA at 65C for Dynabeads) for 30 min, treated with proteinase K for 1 hr at 37C, and de-crosslinked at 65C overnight. DNA fragments were purified with a PCR purification kit (Qiagen). Libraries were prepared using a NEBNext Ultra™ II DNA Library Prep Kit for Illumina (New England Biolabs) as per the manufacturer’s instructions.

All libraries were sequenced on an Illumina NextSeq 500 or 550 platform to obtain 36-37 bp paired-end reads. Two to four biological ChIP-seq replicates were performed in all experiments.

### Hi-C

Hi-C assays were performed as described^22, 23^. The cerebellum from postnatal day 6 or 22 mice or cerebellar vermis from mice at day 1, 2, 4, 7, 8, 14, or 28 after electroporation were homogenized with 1% formaldehyde, followed by quenching with glycine. For the genetic mini-screen, chromatin regulators targeted using dCas9-KRAB mice were subjected to FANS-Hi-C analyses at day 8 after electroporation when significant interactions were formed between regions within the enhancer-dense or gene-dense subcompartment. The rapid knockdown of target genes within ∼3-6 days using dCas9-KRAB^72^ was confirmed using sync-TRAP-seq analyses. Etv1 knockdown using dCas9-KRAB mice was subjected to Hi-C analyses at day 8 or 14 after electroporation, due to increases in Etv1 expression during later stages of granule neuron development. Finally, chromatin regulators targeted using Gabra6-Cre; LSL-Cas9-EGFP mice were subjected to FANS- Hi-C analyses at day 28 after electroporation, due to the ∼1 week needed for Gabra6-Cre expression in developing granule neurons^32, 73^ and the additional ∼2-3 weeks required to deplete target gene expression using wild type Cas9 in postmitotic neurons^22, 74, 75^.

Hi-C libraries were generated and sequenced on the Illumina NextSeq 500 or 550 platform to obtain 36-37bp paired-end reads. Two to six biological Hi-C replicates were performed in all experiments.

### SPRITE

SPRITE experiments were performed as described with modifications^25^. Briefly, the cerebellum from postnatal day 22 mice was fixed with 1% formaldehyde in PBS. ∼10 billion molecules of DNA-protein complexes (∼6000 cells) were coupled with 1.7 ml of NHS-Activated Magnetic Beads (Pierce) per experiment. After five rounds of split-barcoding reactions (DPM, odd, even, odd, and terminal odd), SPRITE libraries were generated and sequenced on the Illumina HiSeq X or NovaSeq platform (Admera Health) to obtain 150bp paired-end reads. Two biological SPRITE replicates were performed.

### DNA FISH

DNA FISH assays were performed as described with modifications^22, 76^. The cerebellum was fixed with 4% PFA and 4% sucrose by transcardiac perfusion. Sagittal cryosections (12 mm thickness) were prepared using a cryostat (Leica Biosystems or Epredia) and cerebellar sections containing the vermis area were used for analyses. Sections were treated with sodium citrate solution (10mM sodium citrate, pH 6.0) for 10 min at 70C and cooled down to room temperature. Sections were dehydrated with sequential treatment of ice-cold 70% EtOH for 2 min, ice-cold 90% EtOH for 2 min, and ice-cold 100% EtOH for 2 min, and air-dried for 5 min. For hybridization, genomic DNA was denatured by incubating glass plates with buffer containing 2x SSC and 70% formamide for 2.5 min at 73C, followed by incubation with 2x SSC and 50% formamide for 1 min. All probes were made from BAC clones (BACPAC Resources Center) including RP24-337G9 (*Ptprt* locus), RP24-209O19 (*Plcb4* locus), RP24-120L14 (*Rims1* locus), RP24-88C18 (*Dpf3* locus), RP24- 252J23 (*Fut8* locus), RP24-153C13 (gene-dense subcompartment locus), and RP24-358P20 (nuclear speckle locus) by nick translation with chemical coupling using an Alexa Fluor succinimidyl ester (AF555 for *Ptprt*; AF647 for *Plcb4*; AF488 for *Rims1*; Alexa647 for *Dpf3*; AF555 for *Fut8*; AF555 for the gene-dense locus; AF647 for the nuclear speckle locus) (Thermo Fisher Scientific). The hybridization solution contained ∼70-80 ng of each labeled probe, 6 mg of mouse Cot-1 DNA (Thermo Fisher Scientific), and 10 mg of sheared salmon-sperm DNA (Thermo Fisher Scientific) in hybridization buffer (10% dextran sulfate, 50% formamide, 2xSSC). The probe mixtures were denatured at 73C for 5 min before use. Denatured sections and probes were sealed and incubated at 37C overnight. On the next day, coverslips were removed and washed once in 2x SSC and 50% formamide solution for 15 min at 45C and three times in 2x SSC for 5 minutes at 45C with gentle agitation. Sections were washed once with PBS and stained with the Hoechst dye. After washing with PBS, sections were mounted with Fluoromount-G mounting medium (Southern Biotech). Images were acquired at the Biological Imaging Facility on a confocal laser scanning microscope (Leica Microsystems, TCS SP8 confocal) using a 63x objective lens (NA 1.40). A stack of 0.3 µm thick optical sections was acquired for each field of view in the UV, red, and far-red channels. Two to three biological DNA FISH replicates were performed in all experiments.

### Immunohistochemistry

Cerebellar sections were prepared and labeled with the relevant antibodies as previously described^32^. A modified heat-induced antigen retrieval protocol^77^ was used for Etv1 staining. Briefly, sections were treated with a sodium citrate solution (10 mM sodium citrate, 0.05% Tween- 20, pH 6.0) under high-pressure for 5 minutes using an Instant Pot® Duo Plus, followed by cooling to room temperature. Secondary antibodies included anti-mouse or rabbit IgG Alexa488 (Thermo Fisher Scientific, A11029 or A11034) or anti-rabbit Alexa647 (Abcam, ab150075).

### DNA FISH combined with immunohistochemistry

Cerebellar sections from 6-8 week-old mice or from mice at day 4, 8, 14, or 28 after electroporation were prepared and treated with sodium citrate solution as described above. Sections were then labeled with the relevant primary and secondary antibodies. After extensive washing of secondary antibodies using PBS, sections were fixed again with 4% PFA in PBS for 10 min at room temperature, washed with PBS for 5 min twice and with 2xSSC once for 5 min, and treated with ice-cold HCl solution (0.1M HCl, 0.7% Triton-X) for 15 min at room temperature. Subsequently, DNA FISH procedures including prehybridization, hybridization, and washing steps, and imaging were performed as described above. Two to three biological replicates were performed in all experiments.

### RNA-seq analyses

For bulk RNA-seq analyses, sequenced reads were aligned to the mm10 reference genome with HISAT2 using the public server at https://usegalaxy.org/ and normalized by library size. Gene annotations were derived using GENCODE version M11.

For sync-TRAP-seq analyses, Smart-seq3 prepared libraries were aligned to the mm10 reference genome using the Smart-seq3 github pipeline (https://github.com/sandberg-lab/Smart-seq3)^70^. Reads from Trio RNA-seq prepared libraries were trimmed 5bp from the 5’ end according to the manufacturer’s instructions (TECAN) and aligned to the mm10 reference genome with HISAT2 using the public server at https://usegalaxy.org/ and normalized by library size. Transcripts considered for analyses were selected based on the presence of H3K4me3 and H3K27ac peaks at the TSS and on higher normalized counts following GFP immunoprecipitation in Rpl10a-GFP electroporated samples compared to non-electroporated control samples. Transcripts enriched at day 4 versus day 8 after electroporation (log_2_ fold change >1 or <-1) and enriched upon Rpl10a- GFP immunoprecipitation compared to total RNA (log_2_ fold change >1) were used to generate modules for visualization on the single nucleus RNA-seq (snRNA-seq) UMAP plot.

snRNA-seq datasets from cerebellar lobules in adult mice with cell type annotations or from the developing cerebellum were obtained^44, 78^ and analyzed using Seurat 4^79^. Briefly, granule neurons or all cells from lobule 4/5 in the adult cerebellum or granule neurons in the P11 developing cerebellum were extracted for downstream analysis. Read counts were normalized and variance stabilized using sctransform and subjected to dimensionality reduction using PCA preprocessing followed by UMAP embedding^79^. Module scores were calculated from the average expression of genes within the module, subtracted by the levels of similarly expressed control genes^79, 80^. Differential gene expression analyses were performed using pair-wise negative binomial tests with edgeR^81^ and the false discovery rate (FDR) was calculated for all genes.

### ChIP-seq analyses

ChIP-seq reads were aligned to the mm10 reference genome with Bowtie2 using the public server at https://usegalaxy.org/ and normalized by library size. Peak calling was performed using MACS2^82^. H3K27ac peaks >2Kb distal to H3K4me3-bound TSSs were considered as enhancers. The ranking of enhancer domains was calculated as previously described^36^. Briefly, H3K27ac peaks within 12.5Kb were stitched together to define enhancer domains. H3K27ac reads within each enhancer domain were normalized by library size, input subtracted, and ordered by rank.

To calculate the average ChIP-seq profile at enhancer domains, the centers of enhancer domains were aligned and the read density was calculated as a function of the distance to center. The standard error of the mean was calculated using two to four biological ChIP-seq replicates. Differential H3K27ac ChIP-seq analyses were performed using pair-wise negative binomial tests with EdgeR and the FDR was calculated for all enhancers.

Transcription factor motif analyses were performed using Homer^83^ to identify known motifs located at ATAC-seq peaks overlapping with H3K27ac-bound enhancers in developing cerebellar granule neurons as described^22, 84^. P-values < 0.05 were converted to negative log_2_ scale for visualization.

### Hi-C analyses

#### Alignment and Juicer analyses

Hi-C reads were aligned to the mm10 reference genome with Bowtie2 and HiC-Pro as described^85^. Uniquely mapped paired-end reads were assigned to MboI restriction fragments and valid pairs with a minimum genomic distance of 1Kb were filtered for PCR duplicates. Contact frequency as a function of genomic distance was calculated as described with modifications^86^. The frequency of valid pairs was calculated as the sum of observed contacts per log_2_ distance bin divided by all observed contacts. Interaction matrices were normalized using Knight-Ruiz (KR) matrix balancing and observed divided by expected (o/e) transformation and visualized using Juicebox^63^. HiCCUPS^23^ was used at 5Kb, 10Kb, and 25Kb resolutions to identify short-distance interactions, and at 25Kb, 50Kb, and 100Kb resolutions to identify ultra-long-distance interactions with o/e frequencies >2.5 and spanning more than 3Mb. Arrowhead was used at 5Kb, 10Kb, and 25Kb resolutions to identify domain boundaries and intersected with short-distance interactions to identify TAD anchors enriched for convergent CTCF binding sites as described^23, 62^. A/B compartment strength was derived from the eigenvector of the Hi-C correlation matrix^23^ using 25Kb or 500Kb bins. Genomic loci enriched for the active histone modification H3K27ac and de-enriched for the repressive histone modification H3K9me3 were assigned a positive eigenvector and denoted as the active A compartment, while genomic loci with a negative eigenvector were denoted as the repressive B compartment. Aggregate peak analysis (APA)^23^ was performed using the indicated groups of ultra-long-distance interactions at 25Kb resolution. The interaction frequency in the central bin was normalized to the mean of the corner bins.

#### Differential interactions with biological replicates

Differential interactions between regions forming ultra-long-distance interactions across cerebellar development were calculated from biological replicates using the edgeR package^81^. Raw interaction counts were normalized by KR balancing and o/e transformation using the offsets parameter, and a generalized linear model-based test was used to identify significant differential interactions.

#### Identification of transcriptionally active inter-chromosomal subcompartments

Dimensionality reduction using Hi-C datasets from P22 cerebellum was performed on the ultra-long-distance interaction matrix between regions in the active A compartment at 500Kb resolution. The intra-chromosomal and inter-chromosomal interaction matrix was normalized by KR balancing and o/e transformation. PCA was computed using the covariance matrix of normalized contacts, and non-negative matrix factorization NMF was computed directly from the normalized contact matrix in Matlab. The NMF rank (k) was approximated by computing at each k the linear regression of lower and higher k residuals and minimizing the joint error sum of squares from the linear regressions^87^. PCA separated the gene-dense and enhancer-dense subcompartments into positive and negative scores of PC2, while NMF separated the gene-dense and enhancer-dense subcompartments into two factors. PCA plots were used to assess the relationship between A compartment strength (PC1) and gene-dense versus enhancer-dense subcompartment scores (PC2).

We observed that NMF factors exhibited modestly higher correlations with identifiable chromatin features compared to PCA analyses. Thus, the top 100 bins at 500Kb resolution with the highest NMF factor scores for the gene-dense subcompartment or the enhancer-dense subcompartment were selected for downstream analyses. To identify at higher resolution the genomic loci associated with transcriptionally active subcompartments, the genome was divided into 5Kb bins and their normalized inter-chromosomal interactions with the top 100 500Kb bins of the gene-dense and enhancer-dense subcompartments were calculated. Genomic loci that belonged to the repressive B compartment were excluded. 5Kb bins with >3-fold enrichment for the gene-dense versus enhancer-dense subcompartment were assigned to the corresponding enriched subcompartment.

#### Aggregate inter-chromosomal interaction analyses

Mean inter-chromosomal contact frequencies were computed between each 25Kb bin of a given genomic locus and all enhancer-dense regions, and visualized as a heatmap aligned to the locus coordinates. To compare multiple loci, the mean contact frequencies between each entire locus and all enhancer-dense or gene-dense subcompartment regions were calculated.

#### A-B compartment interactions

A and B compartment interactions were calculated using the mean normalized long-distance intra-chromosomal interactions (>5Mb) between genomic loci within the enhancer-dense or gene-dense subcompartment and regions within the B compartment at 5Kb resolution. For each chromosome, the top 20 negative eigenvector values at 500Kb resolution were derived from Hi-C datasets of granule neurons at day 8 after electroporation and used to define B compartment regions. The sign of eigenvector values was assigned using H3K27ac and H3K9me3 levels over chromosomes.

#### Local interactions and FIRE analyses

Local interactions were calculated using the mean normalized interactions between all 5Kb bins within each region of the enhancer-dense or gene-dense subcompartment.

FIRE analyses of local *cis*-regulatory interactions were performed using FIREcaller^88^. Briefly, FIREs (p-value < 0.05) were derived from Hi-C datasets of control or Rad21 conditional knockout cerebellum at 40Kb resolution. Differential normalized interactions at FIREs between the control and Rad21 conditional knockout conditions were then calculated from biological replicates using the limma package^89^.

#### PCA of Hi-C datasets from mini-genetic screen

PCA using Hi-C datasets from this study and our previous studies^22^ was performed on the eigenvector of the correlation matrix for datasets at 500Kb resolution. For the eigenvector, genomic loci within the active A compartment were assigned a positive value as described above.

#### Annotation of genomic features

Gene features including gene length, exons, and transcriptional start and termination sites were obtained from GENCODE mouse version M11. Gene ontology analyses for genes overlapping with transcriptionally active subcompartments were performed using DAVID Bioinformatics Resources 6.8^90^ with a background of all expressed genes in the cerebellum.

### DNA FISH and immunohistochemical analyses

For each cell, DNA FISH signals were subjected to automated identification using custom macros in ImageJ as previously described^22, 91^.

Radial positioning of DNA FISH or immunofluorescence signals was analyzed using a single z-section in the middle third (z-axis) of a granule neuron nucleus to ensure a maximal x-y area for calculations. For DNA FISH analyses, the z-section containing the centroid of the DNA FISH probe signal was used. Loci failing to meet these criteria were excluded from analysis. For immunohistochemical analyses, the top 1% pixels by intensity for each neuron were identified. The radial distances from the centroid of the nucleus to the DNA FISH probe signal or immunofluorescence signals were then calculated, where 0 represents the nuclear center and 1 represents the nuclear membrane.

### SPRITE analyses

SPRITE demultiplexing and read count normalization were performed as described with modifications^25^. The DPM sequence was identified from the beginning of read 1 and the odd, even, odd, and terminal odd sequences were identified from read 2 using the Guttman laboratory SPRITE pipeline (https://github.com/GuttmanLab/sprite-pipeline) and together this sequence combination represented the barcode sequence. After trimming the DPM sequence from read 1, the read was aligned to the mm10 reference genome. Mapped reads with MAPQ scores > 30 and a fully identified barcode sequence were kept. Identically mapped reads for a particular barcode sequence were considered PCR duplicates and filtered. After preprocessing, all mapped genomic coordinates with the same barcode sequence were grouped together as a SPRITE cluster. To generate pairwise contact maps, each pairwise interaction was downweighted by 2/n, where n is the total number of reads within a SPRITE cluster. For comparisons with Hi-C analyses, pairwise contacts < 1Kb were removed. Downweighted contacts were then normalized with KR matrix balancing using Juicer^63^. To analyze long-distance interactions, genomic loci within SPRITE clusters were assigned to 500Kb bins. Within each SPRITE cluster, overlapping reads in a bin were counted only once, focusing downstream computations on long-distance genome structure^25, 92^. Clusters containing a minimum of 2 bins and a maximum of 1024 bins were analyzed. For each genomic bin, its observed frequency distribution over a range of cluster sizes was calculated. For a group of genomic bins, the observed frequency distributions were averaged. Genomic loci that did not form ultra-long-distance interactions in Hi-C analyses were used to generate control frequency distributions. SPRITE subclusters were generated based on the genomic distances of loci from either the same chromosome (<5MB or >20Mb, intra-chromosomal contacts) or from different chromosomes (inter-chromosomal contacts) in relation to a reference genomic bin. Reference genomic bins with low coverage, defined as those observed in fewer than 1000 clusters or 100 subclusters, were excluded from frequency distribution analyses.

To calculate interaction frequencies with active subcompartments, SPRITE subclusters harboring regions located on different chromosomes relative to a reference genomic bin were considered. The fraction of bins, excluding the reference genomic bin, that overlapped with regions within the enhancer-dense or gene-dense subcompartment was calculated for each SPRITE subcluster. The mean fractional overlap for each group of reference genomic bins was then computed across a range of SPRITE subcluster sizes and defined as the observed inter-chromosomal interaction frequency. Genomic bins containing regions that did not form ultra-long-distance interactions were used to generate control interaction frequencies for normalization and genomic bins with low coverage were removed as described above.

Dimensionality reduction was performed on the DNA cluster matrix from SPRITE. SPRITE subclusters harboring bins located on different chromosomes with respect to each reference genomic bin were generated. SPRITE subclusters were divided based on size ranges (2, 3-32, or 33-1024 bins) to assess the role of DNA cluster size on genomic interaction preferences. Reference genomic bins observed in less than 200 subclusters were excluded. In a subset of analyses, reference genomic bins overlapping with regions forming ultra-long-distance interactions were selected. A bin ID x subcluster ID matrix was generated with each entry of the matrix containing a 1/n weight, where n represented the size of the subcluster, to ensure that a bin’s weight is proportional to the number of unique bins within the subcluster. Each bin in the matrix was then normalized by the sum of weights for the bin. The bin x subcluster matrix was next subjected to PCA preprocessing followed by UMAP embedding using the Monocle 3 package^93^. When considering UMAP parameters, we noted that inter-chromosomal genomic interactions between any two regions within a nuclear subcompartment only occurred in a few percent of cells, as measured in DNA FISH analyses. Imaging also revealed numerous DNA-protein clusters for each subcompartment within granule neuron nuclei and suggest stochastic and possibly transient association between genomic loci within a subcompartment. In contrast, the chromosome identity of genomic loci strongly influenced their inter-chromosomal interaction preferences due to the packing of chromosomes within territories. To preserve global genome structure, including the sparser interactions between chromosomes, we used high n neighbors (∼20% of bins) and high minimum distance (0.9) for UMAP to also preserve global topological structure for visualization. Louvain community detection was used to identify partitions, which represent groups of bins with similar genomic interaction preferences in UMAP-reduced space. The chromosome mixing index was calculated by determining the chromosome with the maximum number of bins in the partition, dividing this maximum number of bins by the total number of bins in the partition, and then transforming the ratio to a negative natural log scale. A higher chromosome mixing index reflected a broader assortment of chromosomes represented within the partition.

For linear CpG density, the mean CpG density within a 10Mb region surrounding each 100Kb reference genomic bin was calculated. For 3D SPRITE cluster CpG density, we calculated the weighted mean CpG density of regions >3Mb away from each 100Kb reference genomic bin within SPRITE clusters. Contacting regions were downweighted by SPRITE cluster size.

### IGM genome structure-modeling and analysis

We applied our Integrative Genome Modeling (IGM) platform^42^ to generate a population of 1000 single cell 3D genome structures, in which the accumulated genome-wide chromatin contacts across the population are statistically consistent with the Hi-C experiments. We previously demonstrated that structures generated by IGM using Hi-C data can accurately predict with high correlation independent experimental data that map the positions of genomic regions within the nuclear topography. This includes data from SON and Lamin B1 TSA-seq experiments, lamin B1 DamID data, GPseq data, and association frequencies of genomic regions to nuclear speckles, lamina and nucleoli from DNA MERFISH data^42, 43, 94, 95^.

#### Structure representation

The diploid genome is segmented into chromatin regions of 100Kb DNA sequence length, each represented by a spherical domain with radius = 50.6 nm and a genome volume occupancy of 40%, following a procedure as described^42^. Autosomes are represented by two homologous chromatin copies, leading to a total of N = 51,907 chromatin domains in the diploid genome forming a total of 40 chromosome chains. The mouse cerebellum cell nucleus is modelled as an ellipsoid with semi-axes *a* = 3050 nm, *b* = 2350 nm, and *c* = 2350 nm, based on the results of our microscopy measurements.

#### Optimization procedure

The structure optimization is formulated as a maximum likelihood estimation problem using an iterative optimization scheme^42, 94, 96^. Briefly, given the Hi-C based contact probability matrix **A,** we reconstruct the coordinates **X** of all 1000 3D genome structures, together with a latent indicator tensor **W**, which specifies the contacts of homologous genomic regions in each individual structure of the population. We jointly optimize the structure population **X** and the contact tensor 𝐖 by maximizing the log-likelihood of the probability log 𝑃(𝐗|𝐀, 𝐖) = log 𝑃(𝐀, 𝐖|𝐗) following our established population-based modeling approach^42^. Optimizations are initiated with random chromosome configurations.

#### Hi-C data preprocessing

We used the P22 Hi-C data as input information and employed our standard pipeline for preprocessing Hi-C data^42, 43, 94^. This process involved removal of regions with low Hi-C coverage (<3%), matrix balancing through KR normalization conversion of normalized count to contact probabilities, and outlier removal as described in^42^.

#### Prediction of speckle and nucleoli locations in single cell models

We predict the nuclear locations of nuclear speckles and nucleoli in each single cell model as described before^42, 43^. In each single cell structure, a chromatin interaction network (CIN) is calculated for all chromatin regions ***S*** known to have a high propensity to be associated with the nuclear body. To identify this association, we used published SPRITE data^25^ for speckles and published NaD-seq data of F121-9-CASTx129 cells^97^ for nucleoli: each 100Kb bead is marked as nucleoli-associated if its NaD-seq value falls within the top 10 percentile of the genome-wide distribution. Each vertex of CIN represents a 100- Kb chromatin region in ***S***. An edge is established between two vertices *i, j* when their corresponding chromatin regions are in physical contact within the structure. We then employed the Markov Clustering Algorithm (MCL) to identify highly connected subgraphs within the CIN for each genome structure, utilizing the *"mcl"* tool available in the *MCL-edge* software^98^. The positions of each nuclear body within each genome structure are then determined by calculating the geometric centers of all chromatin regions in each subgraph that contain more nodes than a specified threshold. This threshold is set at 3 for speckles, and at the 75-th percentile of cluster sizes for nucleoli. We have previously validated this approach, demonstrating that it produces approximate speckle locations, which reproduce experimental SON-TSA-seq data with high correlation^42, 43^.

#### Identifying spatial partitions of subcompartments as highly connected interaction clusters

To represent individual spatial partitions of the enhancer-dense, gene-dense, and speckle subcompartments, we applied the MCL procedure to identify highly connected subgraphs. CINs were constructed from genomic regions assigned to each subcompartment at 100Kb resolution, and MCL was run as described above. The result is a list of highly connected subgraphs in each structure, where individual subgraphs contain densely connected regions of the same subcompartment.

#### Model assessment by 3D-FISH experiments

To assess the genome structure models, we compared the distributions of radial positions for 5 genomic loci as determined by 3D FISH experiments with the predictions generated by our models (Extended Data Fig. 3h). The 5 loci were part of enhancer-dense, gene-dense, or speckle g-DNA chromatin regions. To measure the radial positions in the models we applied the same procedure employed in the analysis of the 3D FISH imaging data. First, for each single cell structure, we projected the positions of each genomic region onto the X-Y plane. Secondly, we calculated the normalized radial position in the X-Y plane. To account for the elliptical asymmetry of the nucleus, the radial position for a specific genomic region *i* within a given cell is calculated as 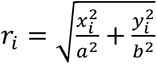, with *a*, *b* as the semi-axis along the X-axis and Y-axis, respectively.

#### Structure feature calculations

Structure features were calculated based on methods as outlined^43^.

#### RAD

The radial position in the nucleus is calculated as the normalized distance relative to the nuclear center at coordinates (0, 0, 0). To account for the elliptical asymmetry of the nucleus, the radial position for a genomic region *i* is calculated as 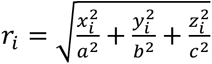, with *a*, *b* and *c* as the semiaxes of the ellipsoid and *x_i_*, *y_i_* and *z_i_* as the coordinates of genomic region *I* in a given genome structure. RAD is the average radial position for a given genomic region over and all homologous copies and structures within the population. <u>RAD-SD</u> is the standard deviation of the mean radial position across the entire population of genome structures.

#### ILF (Interior Localization Frequency)

The interior localization frequency (ILF) of a genomic region *I* is calculated as 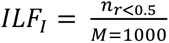, where 𝑛_r<0.5_ is the count of structures in which either one of the two homologous copies of 𝐼 has a radial position lower than 0.5,; 𝑀 is the total number of genome structures in the population.

#### LAF (Lamina Association Frequency)

The lamina association frequency of a genomic region *I* is calculated as 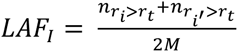, where 𝑀 = 1000 is the number of structures in the population; 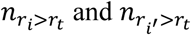 are the number of structures, in which region 𝑖 and its homologous copy 𝑖′ have a radial position *r_i_* larger than the association threshold, 𝑟_t_ = 0.65𝑥𝑅*_nuc_*.

#### SpD (Mean distance to the nearest speckle)

For a genomic region, SpD is the distance to the closest nuclear speckle, averaged over the population of all cells.

SpD-SD: is the standard deviation of SpD across the population of cells.

#### SAF (Speckle association frequency)

The speckle association frequency of a genomic region *I* is calculated as 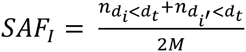, where 𝑀 = 1000 is the number of structures in the population; 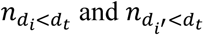 are the number of structures, in which region 𝑖 and its homologous copy 𝑖′ have a bead surface – nearest speckle distance smaller than the threshold, 𝑑_t_ = 500 nm.

#### NuD (mean distance to the nearest nucleolus)

NuD is the distance of a genomic region to the center of the closest nucleolus, averaged over the population of cells. <u>NuD-SD</u> is the standard deviation of the mean nucleolus distance):

#### NAF (Nucleolus association frequency)

The nucleolus association frequency of a genomic region *I* is calculated 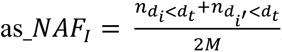, where 𝑀 = 1000 is the number of structures in the population; 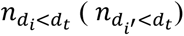 is the number of structures, in which region 𝑖 (its homologous copy 𝑖′ ) has a surface distance to the nearest nucleolus smaller than the threshold 𝑑_t_ = 1000 nm.

#### RG: (mean radius of gyration of local chromatin fiber)

The local compaction of the chromatin fiber at a given locus is calculated by the radius of gyration (RG) for a 500Kb region centered at the locus *i* (i.e. +250Kb up- and 250Kb downstream of *i*), defined as 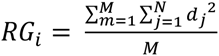, with *N* as the number of chromatin regions in the 500Kb window, and 𝑑_j_is the distance between the chromatin region 𝑗 to the center of mass of the 500Kb region, *M* = 1000 is total number of structures in the population. <u>RG-SD</u> is the standard deviation of the mean RG across the population of genome structures.

#### IPP (mean inter-chromosomal proximity probability)

For each genomic region 𝑖 in each genome structure *m*, we define 𝐸_i_ as the set of genomic regions {𝑗: 𝑗 ≠ 𝑖, 𝑑_ij_ < 500 𝑛𝑚} where 𝑑_!9_ represents the center-to-center distances between *i* and *j* , and is required to be within 500 nm. Within this set 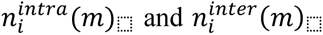 as the number of intra- and inter-chromosomal regions, respectively. IPP is then calculated as: 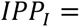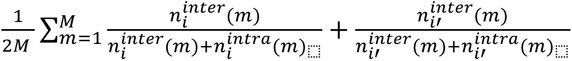 genome structures in the population, and *i* and *i*’ are the two homologous copies of genomic region *I*.

#### TransAB

For a genomic region 𝐼, the trans A/B ratio is defined as 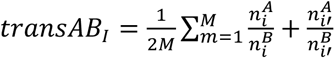 where 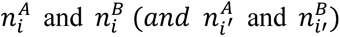 represent the number genomic regions in the A or B compartment in the trans neighborhood set of genomic region *i* (and its homologous copy *i*’) defined as 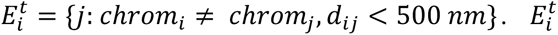 is the set of inter-chromosomal genomic regions located within a distance of 500 nm to genomic region *i*. The mean of the transA/B ratios for a region is then calculated as the average of transA/B ratios from both homologous copies over all structures in the population. The values are rescaled to range between 0 – 1.

#### Fold change analysis of structure features

For each genomic region within each active subcompartment (enh-d, gene-d, speckle), we calculate all 14 structural features. We then calculate the log_2_-fold change in feature values for each subcompartment group by comparing them to values from a corresponding set of randomly selected genomic regions. The reported fold change represents the average value obtained from 100 sets of randomly selected genomic regions.

Visualization of 3D structures was performed with Chimera^99^.

