## Supplemental Information for "Mega-Enhancers Compartmentalize Transcriptionally Active Long Genes in the Brain"

**Supplementary information**

Extended Data

Extended Data Fig. 1 Ultra-long-distance genomic interactions in the developing cerebellum.

**a**, Distribution of intra-chromosomal interactions between genomic regions at various distances, derived using HiCCUPS[1] at 25Kb, 50Kb, and 100Kb resolutions. The median and maximum sizes of all detected interactions from Hi-C analyses of postnatal day 6 (P6) and day 22 (P22) cerebellum are indicated. Ultra-long-distance interactions spanning more than 3Mb significantly increased with cerebellar development. **b**, A/B compartment identity of genomic regions involved in ultra-long-distance interactions. With neuronal maturation, ultra-long-distance interactions were primarily formed between regions in the active A compartment. **c**, A heatmap of differential ultra-long-distance interactions within the A compartment during cerebellar development (false discovery rate (*FDR*) < 0.05, two-sided *P* value from negative binomial distribution with Benjamini–Hochberg post hoc test, n = 3 biological replicates, $\log_2$ fold change and mean centered). **d**, UCSC genome browser tracks of the active histone marks H3K4me3 and H3K27ac at two example regions on chromosomes 1 and 12, which form a subset of the ultra-long-distance genomic interactions shown in Fig. 1c. Abundantly expressed isoforms of *Dst* and *Nrxn3*, which are also marked by H3K4me3 at transcriptional start sites (TSS) in granule neurons, are depicted schematically. **e**, The dimensionality reduction approaches, including principal component analysis (PCA) and non-negative matrix factorization (NMF), used to identify distinct subcompartments and their associated chromatin features. **f**, Regions in the enhancer-dense (red) and gene-dense (blue) subcompartment identified by NMF (Methods) projected onto the PCA plot in Fig. 1d. These NMF-derived subcompartments separate along principal component 2 (PC2) of the PCA plot. **g**, Comparisons of gene, enhancer, or exon density, as well as gene length, among transcriptionally active subcompartments or the entire mouse genome at 500Kb resolution. The enhancer-dense subcompartment was characterized by a high density of enhancers and was enriched for genes spanning over 100Kb (**FDR* < 0.05, ****FDR* < 0.001, Kruskal-Wallis test with Dunn's post hoc test, *n* = 100, 100, 25646 regions for enh-dense, gene-dense, all genome). In contrast, the gene-dense subcompartment was characterized by a high density of genes and exons. Box plots in **g** show median, quartiles (box), and range (whiskers).

Extended Data Fig. 2 Features of the enhancer-dense and gene-dense subcompartments.

**a,** The length of the top 1% of enhancers enriched for H3K27ac in P6 or P22 cerebellum. An increased number of top 1% enhancers that span over 100Kb was observed during cerebellar development ($P = 5.9 \times 10^{-25}$, two-sided Mann-Whitney-Wilcoxon test, *n* = 174, 195 enhancers for P6, P22). **b,** The length of the top 1% of enhancers enriched for H3K27ac in human tissues or cells[2]. Top 1% enhancers were typically longer in human brain regions (magenta) compared to in other tissues (black). **c,** UMAP visualization of snRNA-seq data from lobule 4/5 in the adult cerebellum[3]. Annotations of cerebellar cell types (left), Seurat[4] module scores of genes overlapping with the top 1% of enhancers enriched for H3K27ac (middle), and module scores of genes found within the gene-dense subcompartment (right) are shown. **d,** Density of housekeeping or metabolic genes found within various transcriptionally active subcompartments in the cerebellum, normalized to that found in all other A compartment regions. enh-d: enhancer-dense subcompartment; gene-d: gene-dense subcompartment; speckle: nuclear speckle-associated regions. **e,** Inter-chromosomal interaction frequencies between enhancer domains of various lengths, categorized by intragenic versus intergenic enhancers, and the enhancer-dense subcompartment.

Box plots in **a** and **b** show median, quartiles (box), and range (whiskers).

Extended Data Fig. 3 Higher-order assembly of active subcompartments.

**a**, The normalized frequencies of the *Rims1* locus appearing in SPRITE subclusters selected based on the genomic distance of loci from the *Rims1* locus. Subclusters were classified as being from the same chromosome (<5Mb or >20Mb, intra-chromosomal) or from different chromosomes (inter-chromosomal). **b**, UMAP plots representing the inter-chromosomal interaction preferences of genomic regions forming ultra-long-distance interactions in Hi-C analyses across various ranges of SPRITE subcluster sizes. Top, color-coded chromosomes. Bottom, the enhancer-dense (red) or gene-dense (blue) subcompartment is highlighted. Enhancer-dense regions cluster together in UMAP space (circled in red) within SPRITE subclusters of size 2, and to some extent sizes 3 to 32. **c**, Inset, color-coded partitions obtained from Louvain clustering of UMAP reductions on SPRITE subclusters of size 2 from **b**. Main plot, the correlation between the fraction of enhancer-dense regions within partitions and the degree of chromosome mixing within partitions. Enhancer-dense regions located on different chromosomes were grouped together in UMAP space for SPRITE subclusters of size 2. **d**, Fold change enrichment for each of the 14 structural features (Methods) for enhancer-dense (top), gene-dense (middle), and nuclear speckle-associated regions (bottom). **e**, Violin plots showing the distributions of the mean radial positions (RAD, top left), inter-chromosomal proximity probability (IPP, top right), mean distances to the nearest nuclear speckle (SpD, bottom right), and mean distances to the nearest nucleolus (NuD, bottom left) for enhancer-dense (red), gene-dense (blue), and nuclear speckle-associated regions (gold) (***$P <$

0.001, Mann-Whitney-Wilcoxon test). **f**, t-distributed stochastic neighbor embedding (t-SNE) to visualize the high-dimensional nuclear microenvironment of enhancer-dense (red), gene-dense (blue), and nuclear speckle-associated regions (gold). The nuclear microenvironment of each genomic locus was defined by a high-dimensional feature vector consisting of the ordered list of 14 features that served as topographic markers and are listed in **d**. **g**, The radial distributions of DNA FISH probes targeting the enhancer-dense regions *Rims1*, *Plcb4*, and *Ptprt* (top) and immunofluorescence signals for Sf3a66 and H3K27ac (bottom) as in Fig. 3h. **h**, Comparison of radial positions for selected genomic loci associated with the enhancer-dense subcompartment (red), the gene-dense subcompartment (blue), or nuclear speckles (gold), as predicted from IGM or determined using DNA FISH ($FDR > 0.1$, Mann-Whitney-Wilcoxon test with Benjamini-Hochberg correction, $n$ = 87-113, 2000-4000 loci for DNA FISH, IGM).

Data in **a** show mean and shading denote s.e.m. Box plots in **h** show median, quartiles (box), and range (whiskers).

Extended Data Fig. 4 Changes in transcription and genome organization at various stages of granule neuron development.

**a**, *Grin2b* (left) or *Grin2c* (right) expression in developing granule neurons visualized with UMAP as in Fig. 4b. **b**, The fluorescence intensities and percentages of sorted cells with FANS, as in Fig. 4c, using the cerebellum of a non-electroporated P15 mouse. No cells were detected in the Q2 gate in the non-electroporated control condition (red outline). **c**, Hi-C analyses of intra-chromosomal interactions between the enhancer-dense or gene-dense subcompartment, which belong to the A compartment, and the B compartment during granule neuron maturation *in vivo*. Interactions between the A and B compartments were weakened during granule neuron maturation, with some interactions between the gene-dense subcompartment and the B compartment persisting in adult animals (*$P < 0.05$, **$P < 0.01$, ***$P < 0.001$, Friedman test with Nemenyi's post hoc test, $n$ = 187, 234 loci for enh-d, gene-d). IVE: *in vivo* electroporation. **d**, Hi-C analyses of local interactions within regions of the enhancer-dense or gene-dense subcompartment during granule neuron maturation *in vivo*. Local interactions within regions of the enhancer-dense subcompartment increased during granule neuron maturation (*$P < 0.05$, ***$P < 0.001$, Friedman test with Nemenyi's post hoc test, $n$ = 192, 249 loci for enh-d, gene-d). **e**, The strength of inter-chromosomal interactions between regions within gene-dense subcompartment increased during cerebellar development (left, ***$P < 0.001$, Wilcoxon signed-rank test, $n$ = 261 loci for gene-d) or during various stages of granule neuron maturation *in vivo* (right, **$P < 0.01$, ***$P < 0.001$, Friedman test with Nemenyi's post hoc test, $n$ = 249 loci for gene-d). **f**, Genes and CpG-islands located along the linear genome surrounding enhancer-dense regions at the *Rims1* or *Dpf3* locus. **g**, The average CpG density of the surrounding 10Mb region along the linear genome for all genomic loci. Genomic loci are sorted by linear CpG density, and loci in enhancer-dense subcompartment from Fig. 4j are highlighted in red. **h**, The weighted mean CpG density of regions located >3Mb away from each given genomic locus within 3D SPRITE clusters. Values are centered over the mean of all genomic loci. Genomic loci are sorted by normalized CpG density and loci in enhancer-dense subcompartment from Fig. 4j are highlighted in red. Enhancer-dense regions more peripherally located within the nucleus formed 3D ultra-long-distance interactions with regions of relatively lower CpG density. **i**, Radial positions of genomic loci in the A compartment (magenta) or B compartment (green), as calculated from IGM, compared to the weighted mean CpG density of regions forming ultra-long-distance interactions with these loci in 3D SPRITE clusters. Values are centered over the mean of all genomic loci. The nuclear periphery (peri) and nuclear interior (int) are denoted. Genomic loci in the A compartment interacting with CpG-poor regions in SPRITE clusters tended to be positioned closer to the nuclear periphery.

Box plots in **c**-**e** show median, quartiles (box), and range (whiskers).

Extended Data Fig. 5 Effects of genetic perturbations targeting nuclear proteins on genome architecture and gene expression *in vivo*.

**a**, The sgRNA targeting strategy for knockdown of genes in dCas9-KRAB mice. Two sgRNAs targeting the 100bp region surrounding the TSS were designed for each gene. **b**, Fold change in mRNA levels upon knockdown (KD) or conditional knockout (cKO) of targeted genes compared to their respective control condition ($n$ = 2-11 biological replicates). **c**, Fold changes in inter-chromosomal interactions between regions within the gene-dense subcompartment upon knockdown or conditional knockout of 13 nuclear proteins, as shown in Fig. 5c ($n$ = 2-6 biological replicates). **d**, Hi-C contact maps of a 70Mb region on chromosome 2 in control (top) or Top2b knockdown (bottom) granule neurons. Red indicated interactions between A or B compartment regions, while blue indicated the absence of interactions between the A and B compartments. **e**, Relative frequencies of Hi-C eigenvector scores in control (ctrl) or Top2b knockdown granule neurons *in vivo*. The arrow indicates A compartment regions with high positive eigenvector scores, which are reduced following Top2b knockdown. **f**, Hi-C analyses of ultra-long-distance intra-chromosomal interactions between the enhancer-dense (enh-d) or gene-dense (gene-d) subcompartment and the B compartment in control (ctrl) or Top2b knockdown granule neurons *in vivo*. Top2b knockdown increased interactions between active subcompartments and the B compartment (***$P < 0.001$, Wilcoxon signed-rank test, $n$ = 190, 246 loci for enh-d, gene-d). **g**, The effects of knockdown or conditional knockout of 13 nuclear proteins on inter-chromosomal interactions between regions within either the enhancer-dense subcompartment (top) or gene-dense subcompartment (bottom) compared to their effects on intra-chromosomal interactions between these subcompartments and the B compartment. Among the genetic perturbations, Top2b knockdown led to the strongest increases in interactions between active subcompartments and the B compartment. **h**, Comparison of the fold change in gene expression upon knockdown of Top1/2b with respect to gene length. Top1 and Top2b depletion had little or no effect on the overall expression of neuronal long genes in granule neurons *in vivo*. **i**, Hi-C contact map of a TAD surrounding the neuronal long gene *Rims1* on chromosome 1 in Rad21 conditional knockout compared to control granule neurons. Local interactions within the TAD were reduced upon conditional knockout of Rad21. Local frequently interacting regions (FIREs)[5] identified in control granule neurons are indicated in magenta. **j**, The fraction of FIREs that were significantly upregulated or downregulated upon conditional knockout of Rad21 compared to the control condition ($FDR < 0.05$, two-sided $P$ value from negative binomial distribution with Benjamini–Hochberg post hoc test, $n = 4$ biological replicates). Depletion of Rad21 led to the overall weakening of FIREs. **k**, The fraction of regions within the enhancer-dense or gene-dense subcompartment overlapping with Rad21 conditional knockout-downregulated FIREs. Depletion of Rad21 selectively weakened local interactions within regions of the enhancer-dense subcompartment. **l**, Hi-C analyses of local interactions within regions of the enhancer-dense or gene-dense subcompartment in control or Rad21 conditional knockout (cKO) granule neurons *in vivo*. Rad21 depletion reduced local interactions within regions of active subcompartments (***$P < 0.001$, Wilcoxon signed-rank test, $n = 195$, 261 loci for enh-d, gene-d). **m**, The effects of knockdown or conditional knockout of 13 nuclear proteins on inter-chromosomal interactions between regions within the enhancer-dense subcompartment (top) or gene-dense subcompartment (bottom) compared to their effects on local interactions within regions of these subcompartments. Among the genetic perturbations, Rad21 conditional knockout led to the strongest reductions in local interactions within regions of active subcompartments.

Box plots in **f** and **l** show median, quartiles (box), and range (whiskers). Data in **b**, **c**, **e**, and **h** show mean and error bars or shading denote s.e.m.

Extended Data Fig. 6 Etv1 orchestrates transcription and genome organization in mature granule neurons.

**a**, An image of the cerebellar cortex from P10 mice labeled using the Etv1 (green) and calbindin (red) antibodies together with the Hoechst DNA dye (blue). External granule layer (EGL), molecular layer (ML), Purkinje cell layer (PCL), and internal granule layer (IGL) are denoted. Etv1 protein was selectively expressed in mature granule neurons in the IGL. **b**, Etv1 knockdown or control granule neurons at day 8 after electroporation were isolated using FANS and subjected to RNA-seq analyses. All differentially expressed transcripts (left) or differentially expressed long transcripts spanning over 100Kb (right) are shown ($FDR < 0.01$, two-sided $P$ value from negative binomial distribution with Benjamini–Hochberg post hoc test, $n = 6$ biological replicates). **c**, Enhancers with differential H3K27ac levels upon Etv1 knockdown ($FDR < 0.05$, two-sided $P$ value from negative binomial distribution with Benjamini–Hochberg post hoc test, $n = 3$ biological replicates). **d**, Changes in H3K27ac levels upon Etv1 knockdown at enhancers associated with Etv1 knockdown-downregulated or upregulated genes (***$P < 0.001$, Wilcoxon signed-rank test, $n$ = 168-169 loci). **e**, Fractional overlap of Etv1 knockdown-upregulated or downregulated transcripts ($log_2$ fold change >0.585 or < -0.585) with enhancer-dense, gene-dense, or nuclear speckle-associated regions normalized to the overlap of unchanged genes with these subcompartments. Etv1 knockdown-downregulated transcripts were enriched within the enhancer-dense subcompartment compared to unchanged genes (****FDR* < 0.0001, Fisher's exact test with Benjamini-Hochberg correction). **f**, Hi-C analyses of inter-chromosomal interactions between Etv1 knockdown-downregulated long genes ($\log_2$ fold change < -0.585) and the enhancer-dense or gene-dense subcompartment in control or Etv1 knockdown granule neurons at day 8 or 14 after electroporation. Long genes expressed in an Etv1-dependent manner strengthened their interactions with the enhancer-dense subcompartment during granule neuron maturation *in vivo*, but these developmental changes in genome architecture were blocked by Etv1 knockdown (***P* < 0.01, ****P* < 0.001, Friedman test with Nemenyi's post hoc test, *n* = 105 transcripts). **g**, Hi-C analyses of inter-chromosomal interactions between Etv1 knockdown-upregulated long genes ($\log_2$ fold change > 0.585) and the enhancer-dense subcompartment in control or Etv1 knockdown granule neurons at day 14 after electroporation. Etv1 depletion had little or no effect on the interactions between genomic loci containing Etv1 knockdown-upregulated long genes and the enhancer-dense subcompartment (*P* = 0.67, Friedman test with Nemenyi's post hoc test, *n* = 334 transcripts).

Box plots in **d**, **f**, and **g** show median, quartiles (box), and range (whiskers).

Extended Data Fig. 7 Effects of Brd4 and Ubash3b on transcription and the enhancer-dense subcompartment.

**a**, Brd4 knockdown or control granule neurons at day 8 after electroporation were subjected to sync-TRAP-seq analyses. Differentially expressed transcripts are shown (*FDR* < 0.01, two-sided *P* value from negative binomial distribution with Benjamini–Hochberg post hoc test, *n* = 7-11 biological replicates). **b**, Overlap of Brd4 knockdown-upregulated or downregulated transcripts with enhancer-dense, gene-dense, or nuclear speckle-associated regions normalized to the overlap of unchanged genes with these subcompartments. Brd4 knockdown-downregulated transcripts were enriched within the enhancer-dense subcompartment compared to unchanged genes (***$FDR < 0.0001$, Fisher's exact test with Benjamini-Hochberg correction). **c**, Inter-chromosomal interaction frequencies between regions within the enhancer-dense subcompartment in control or *Ubash3b* enhancer inactivated granule neurons at day 14 after electroporation. *Ubash3b* enhancer inactivation had little or no effect overall on the enhancer-dense subcompartment ($P = 0.42$, Wilcoxon signed-rank test, $n = 195$ loci).

Box plots in **c** show median, quartiles (box), and range (whiskers).

Supplementary Tables

Supplementary Table 1. Sample information for Hi-C or SPRITE analyses.

Supplementary Table 2. Transcripts enriched in granule neurons at day 4 or day 8 after electroporation.

Supplementary Table 3. Differentially expressed transcripts with Top1/2b, Etv1, or Brd4 knockdown in granule neurons.

Supplementary Table 4. Mouse lines and samples used in this study.

Supplementary Table 5. Sequences of sgRNAs for the *in vivo* genetic mini-screen.

### Extended Data Fig. 1

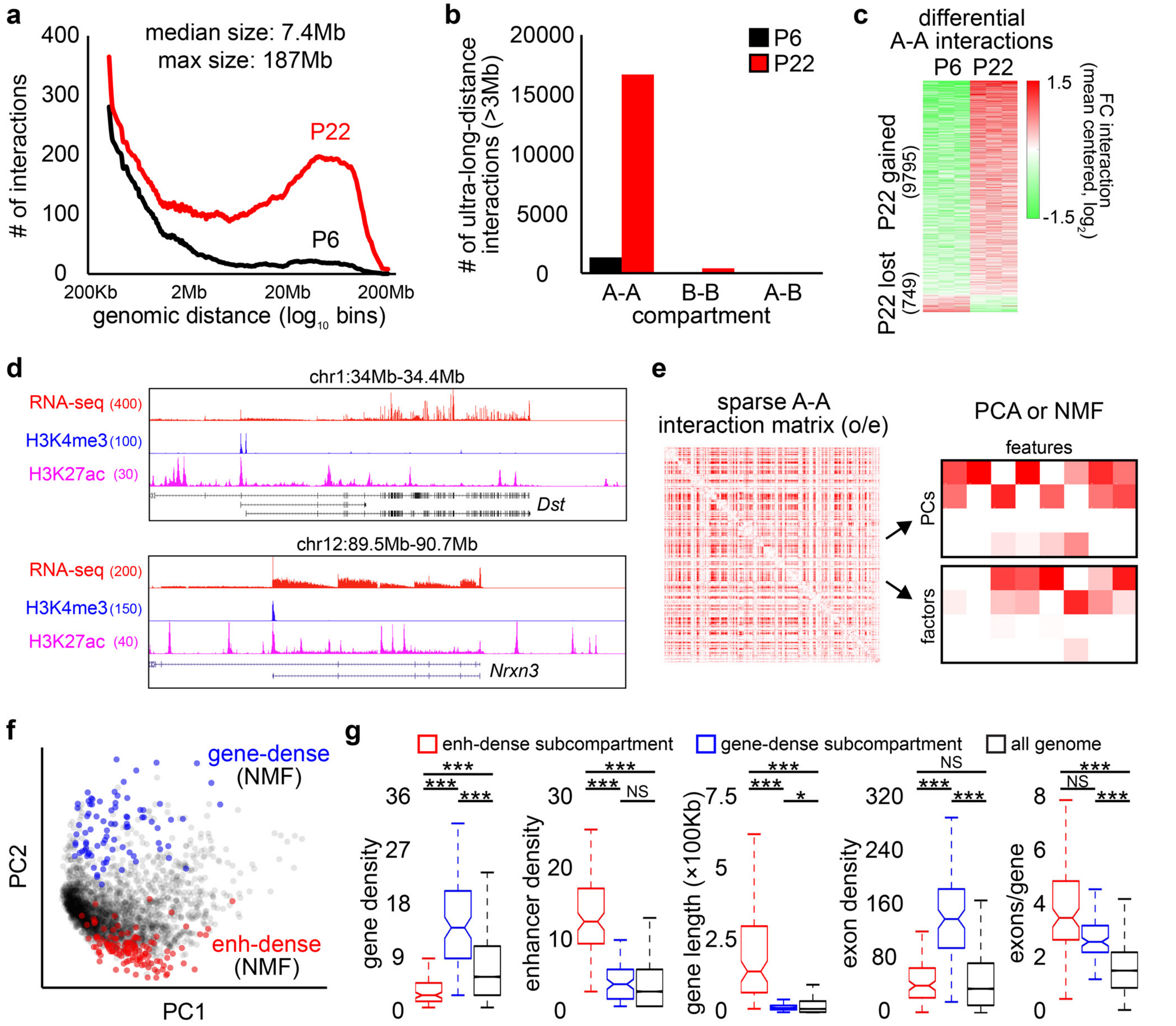

**Extended Data Fig. 2**

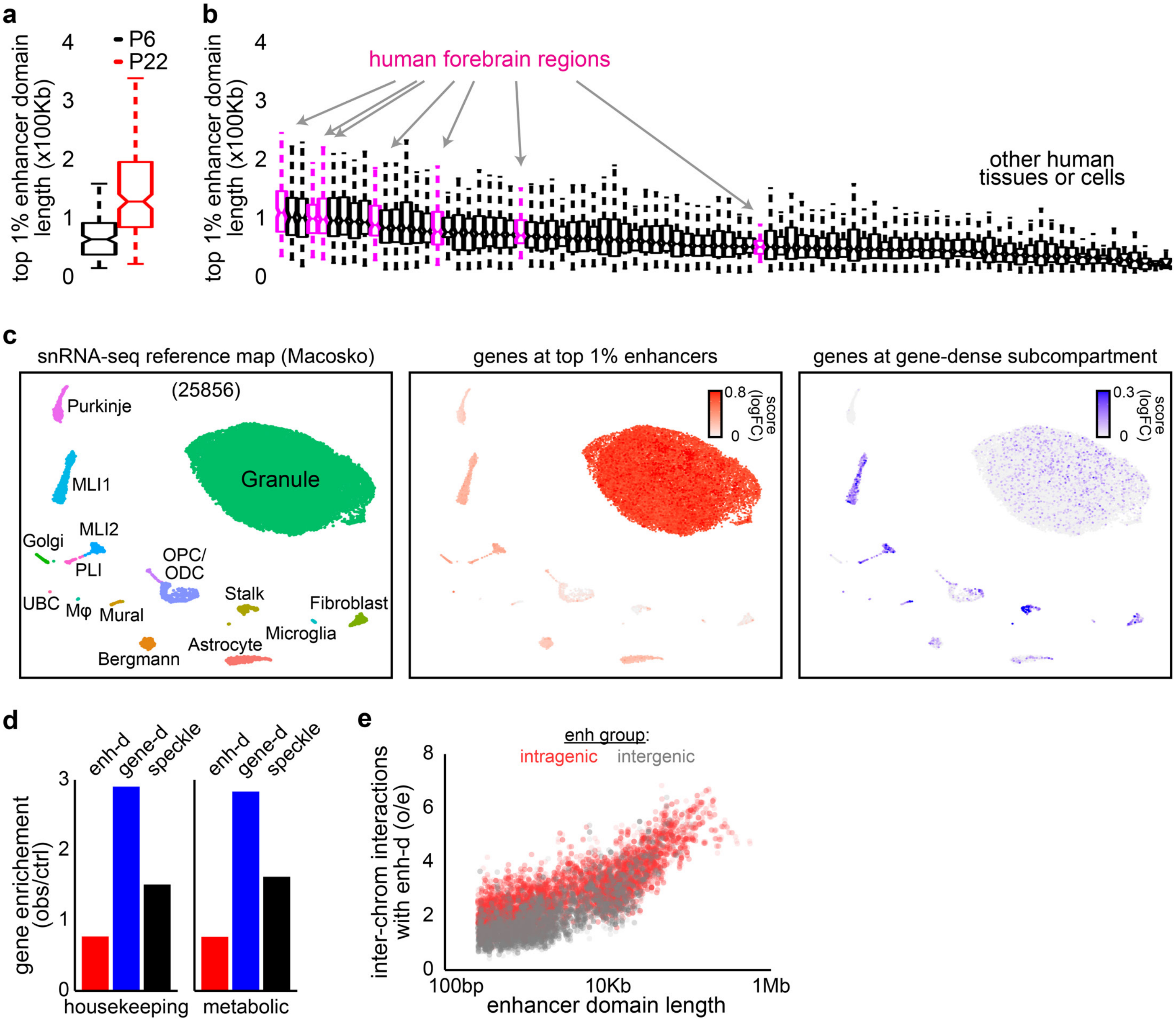

#### Extended Data Fig. 3

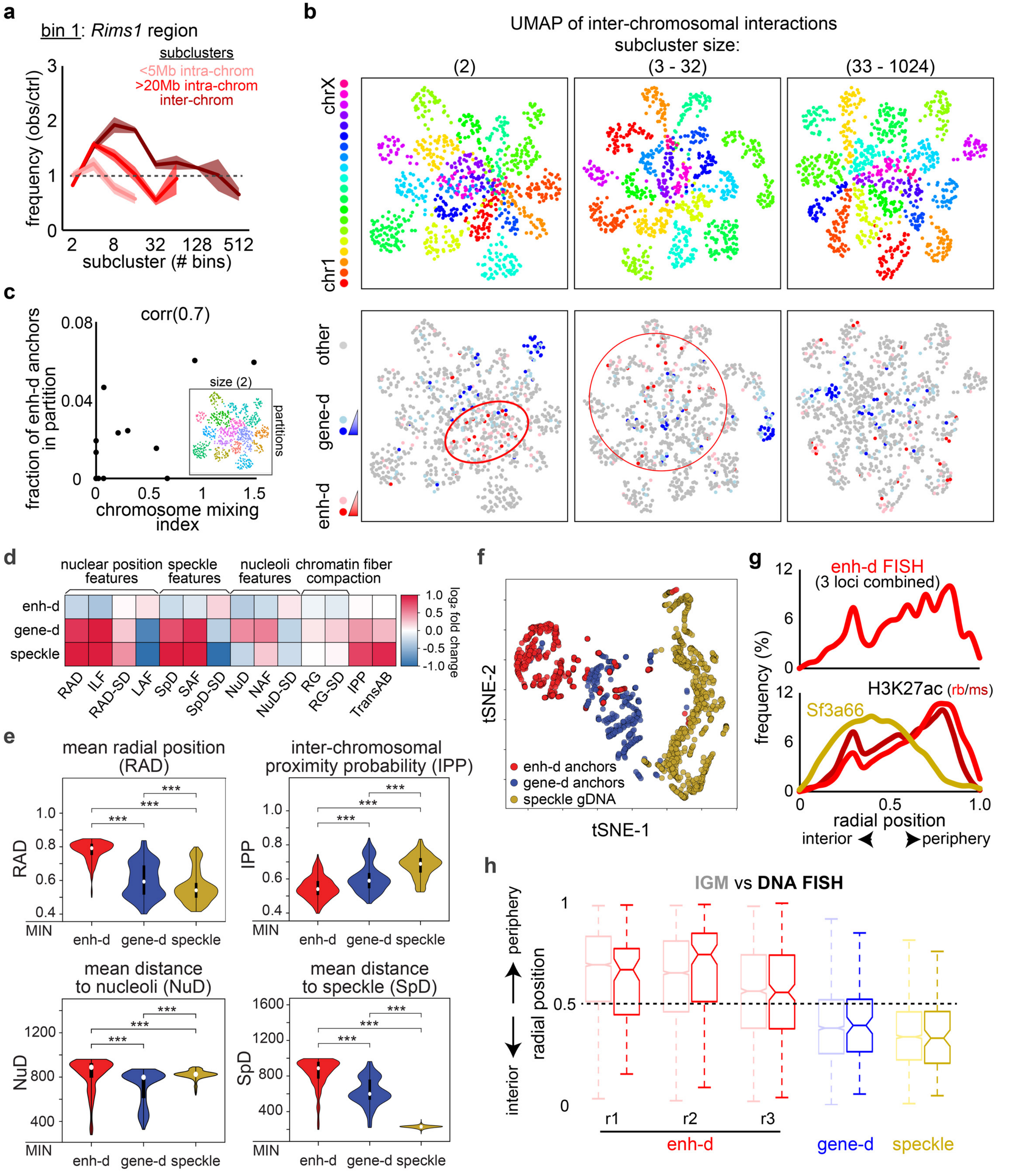

#### Extended Data Fig. 4

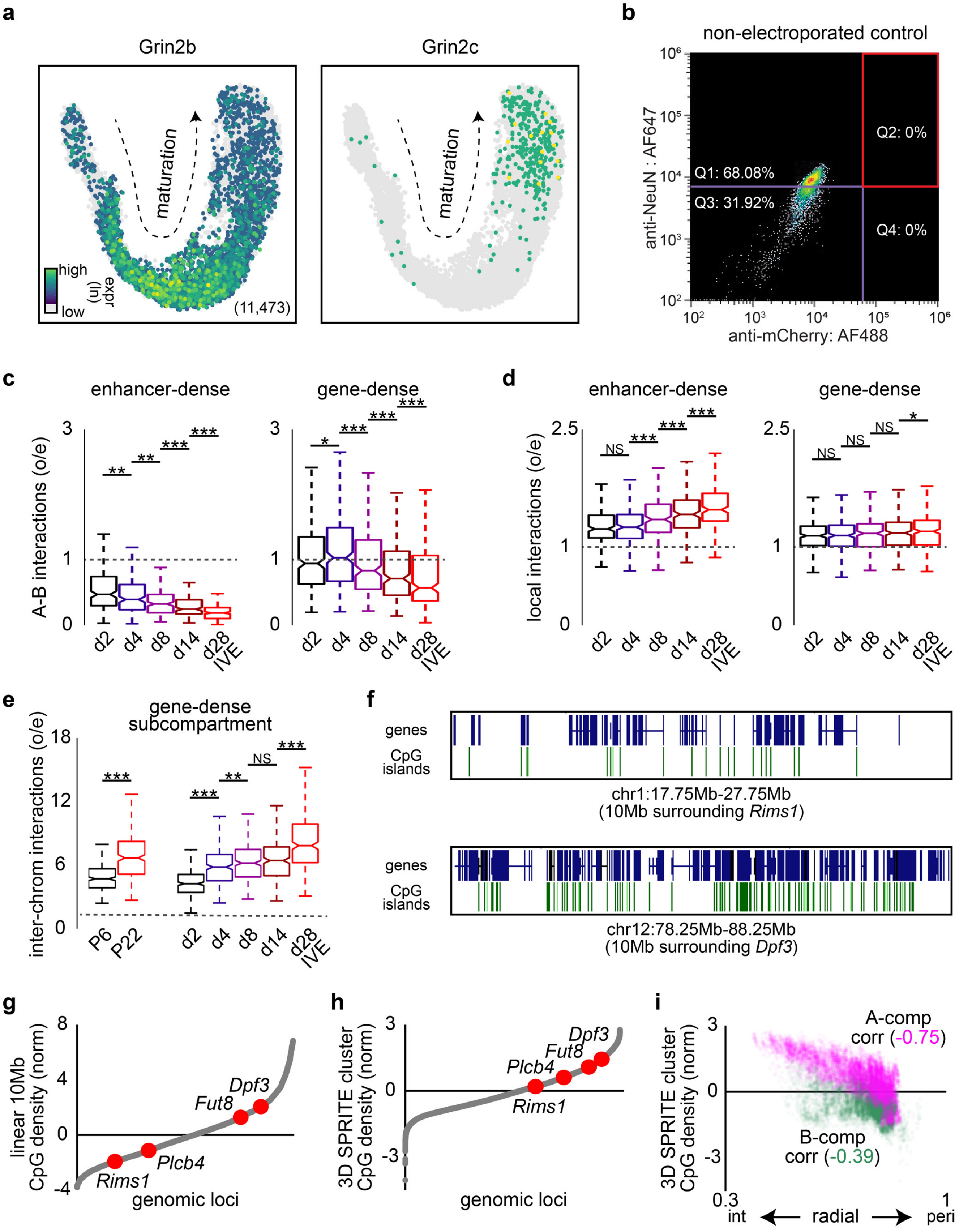

#### Extended Data Fig. 5

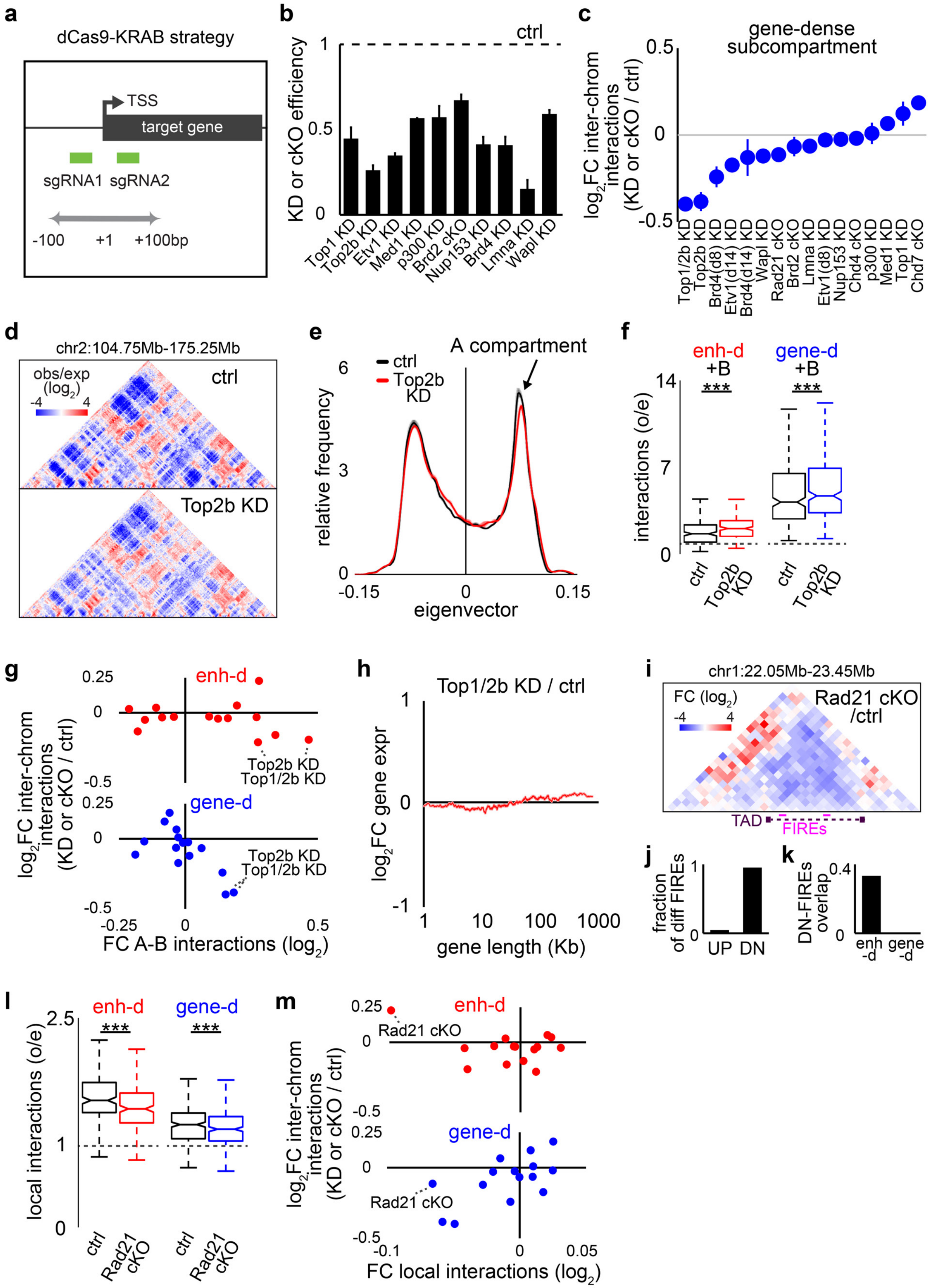

**Extended Data Fig. 6**

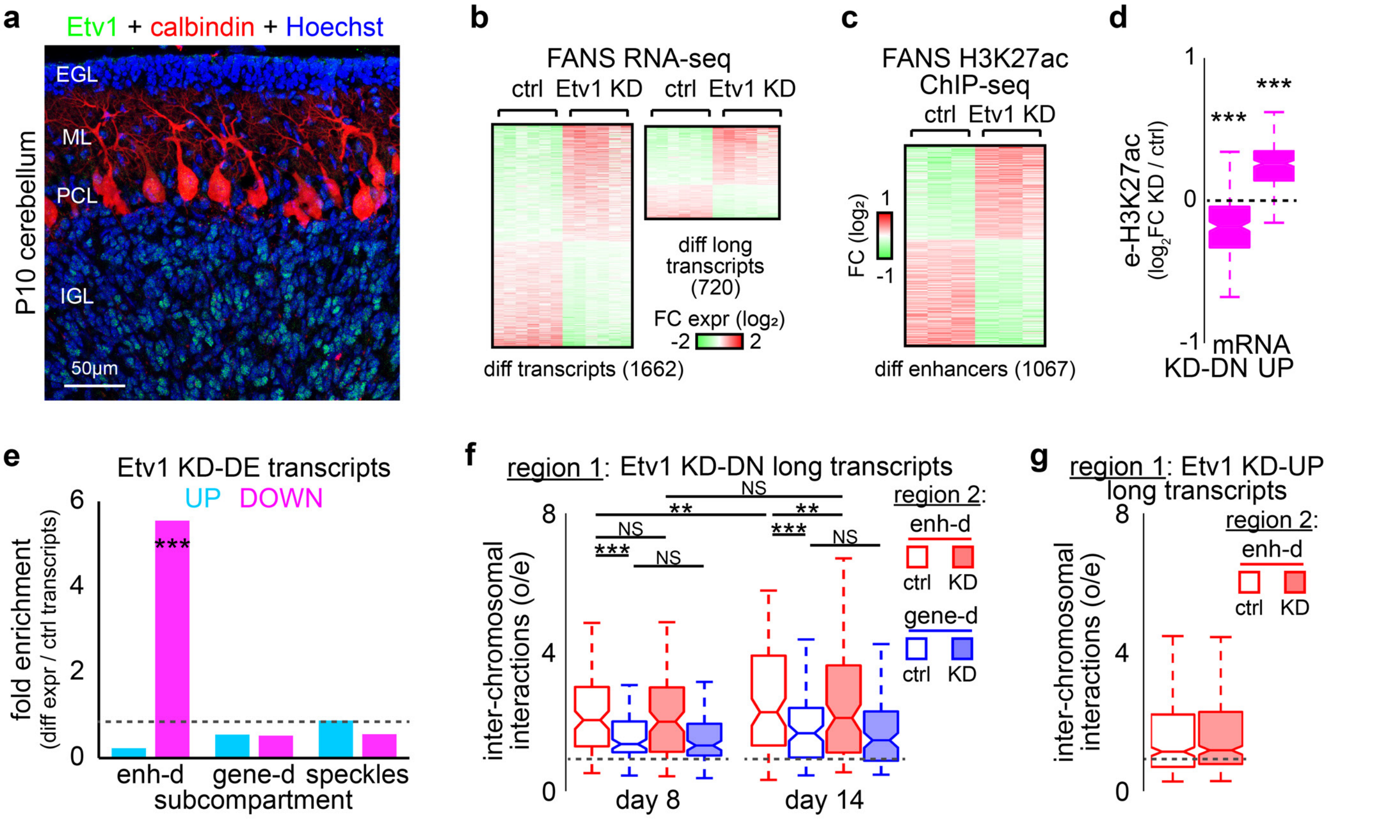

#### Extended Data Fig. 7

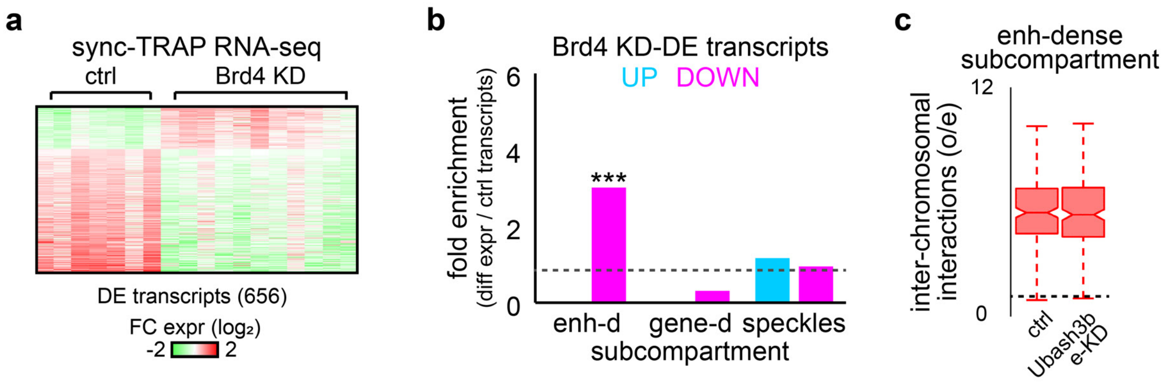
